# Expanding the human gut microbiome atlas of Africa

**DOI:** 10.1101/2024.03.13.584859

**Authors:** Dylan G Maghini, Ovokeraye H Oduaran, Jakob Wirbel, Luicer A Ingasia Olubayo, Natalie Smyth, Theophilous Mathema, Carl W Belger, Godfred Agongo, Palwendé R Boua, Solomon SR Choma, F Xavier Gómez-Olivé, Isaac Kisiangani, Given R Mashaba, Lisa Micklesfield, Shukri F Mohamed, Engelbert A Nonterah, Shane Norris, Hermann Sorgho, Stephen Tollman, Floidy Wafawanaka, Furahini Tluway, Michèle Ramsay, Ami S Bhatt, Scott Hazelhurst

## Abstract

Population studies are crucial in understanding the complex interplay between the gut microbiome and geographical, lifestyle, genetic, and environmental factors. However, populations from low- and middle-income countries, which represent ∼84% of the world population, have been excluded from large-scale gut microbiome research. Here, we present the AWI-Gen 2 Microbiome Project, a cross-sectional gut microbiome study sampling 1,803 women from Burkina Faso, Ghana, Kenya, and South Africa. By intensively engaging with communities that range from rural and horticultural to urban informal settlements and post-industrial, we capture population diversity that represents a far greater breadth of the world’s population. Using shotgun metagenomic sequencing, we find that study site explains substantially more microbial variation than disease status. We identify taxa with strong geographic and lifestyle associations, including loss of *Treponema* and *Cryptobacteroides* species and gain of *Bifidobacterium* species in urban populations. We uncover a wealth of prokaryotic and viral novelty, including 1,005 new bacterial metagenome-assembled genomes, and identify phylogeography signatures in *Treponema succinifaciens*. Finally, we find a microbiome signature of HIV infection that is defined by several taxa not previously associated with HIV, including *Dysosmobacter welbionis* and *Enterocloster sp.* This study represents the largest population-representative survey of gut metagenomes of African individuals to date, and paired with extensive clinical biomarkers, demographic data, and lifestyle information, provides extensive opportunity for microbiome-related discovery and research.

## Introduction

Large population studies can identify lifestyle, genetic and environmental factors that drive gut microbiome composition. Indeed, early population studies established baseline human gut measurements: for example, the MetaHIT study sampled from 124 Europeans catalogued millions of bacterial proteins and identified core gut bacterial functions^1^, and the Human Microbiome Project (HMP) sampled from 242 individuals from the United States, and found high individuality of microbiome composition and conserved core metabolic pathways across microbiomes^2^. More recently, population studies have related the gut microbiome to disease and lifestyle factors: HMP2 identified taxonomic associations with inflammatory bowel disease^3^ and type 2 diabetes biomarkers^4^ in cohorts of 132 and 106 individuals, respectively, and the Dutch Microbiome Project found a strong effect of co-housing on microbiome composition and uncovered several associations between diet, socioeconomics, and environmental exposures and the gut microbiome in a cohort of 8,208 individuals^5^. These studies and others (**STable 1**) have built a strong foundation for the microbiome field. However, as these studies typically focus on high-income populations with relatively homogeneous resource access and disease profiles, their results often do not translate to those living in other geographic areas or regions with more limited resource access, as well as different lifestyle practices, challenges to health and healthcare, and environmental exposures. Additionally, the vast majority of large microbiome studies have relied on a non-random and facility-based convenience-sampling model, which generalizes less well to the population level than more time- and resource-intensive cross-sectional sampling approaches. A notable exception is the FINRISK study, which has highlighted the utility of cross-sectional sampling to study microbiome signatures of mortality risk in a population-representative Finnish cohort^6^.

Despite the high diversity of lifestyle, diet, environment, disease burden, and resource access in low- and middle-income countries (LMICs), which together are home to ∼84% of the world’s population^7^, people living in LMICs are extremely underrepresented in gut microbiome research^8–10^. An excellent collection of targeted studies in focused LMIC communities has found microbiome associations with infectious disease^11–13^, described compositional differences between microbiomes of specific LMIC and high-income country (HIC) populations^14–18^, including in the context of non-communicable disease^19,20^, and shown that microbiome composition can change with migration from LMICs to HICs^21^. However, to more comprehensively capture global lifestyle diversity that affects gut microbiome composition, it is necessary to study more diverse populations living in LMICs, not just those living in extremes of lifestyle (e.g. hunter gatherers) or focused ethnic communities (e.g. the Hmong/Karen people of Thailand). This can be accomplished by designing and executing large, population-representative studies that encompass a range of lifestyles, geography, and disease burden, and which collect high-quality data about these factors. Importantly, it is essential to work within local frameworks that support representative measurements from populations, facilitate input and leadership from local stakeholders, and clearly identify community needs^22–26^.

The Africa Wits-INDEPTH Partnership for Genomic Studies (AWI-Gen)^27^ provides a powerful and versatile framework for broad, multi-country, population-representative and community-engaged microbiome research (**Figure 1**). Nested within the Human Heredity and Health in Africa Consortium (H3Africa), AWI-Gen seeks to identify genomic and environmental factors affecting the changing disease burden among older adults in six African communities in four countries. The study is a partnership between the University of the Witwatersrand and the International Network for the Demographic Evaluation of Populations and Their Health (INDEPTH), a global network of health and demographic surveillance systems (HDSSs) that monitor population health in LMICs. AWI-Gen includes five HDSSs and the Developmental Pathways for Health Research Unit (DPHRU) in Soweto, South Africa, enabling a robust, multicentre research platform. First, HDSSs enable random cross-sectional sampling of individuals across their catchment areas. By contrast, most extant microbiome and genomics data in the public sphere are based on highly non-random recruitment that relies on convenience sampling, where self-selecting participants learn about the study through advertisements or word-of-mouth, which is not optimal for capturing true population-level trends. Second, each HDSS has operated among host communities for over a decade, hiring local field workers and supporting teams that conduct extensive community engagement prior to study approval and through study conclusion^28,29^. This supports improved study design, high participant response and retention, and an assurance that study questions focus on community needs. Finally, AWI-Gen has collected blood and urine biomarkers, captured extensive participant data — demographic, health history, environment, and lifestyle — and genotyped all participants on the 2.2m SNP H3Africa Custom Array^30^. Importantly, AWI-Gen places heavy emphasis on genomic research capacity-building and equitable scientific collaborations. Altogether, AWI-Gen presents a unique opportunity for microbiome research in understudied populations and holds immense potential for associating the microbiome with a rich set of genotype and phenotype data. The first phase of AWI-Gen was conducted from 2012 - 2017, during which we carried out pilot microbiome projects at the Agincourt HDSS and Soweto DPHRU in South Africa^31,32^. The second phase of AWI-Gen, including our microbiome study, was conducted from 2018-2023.

**Figure 1.**
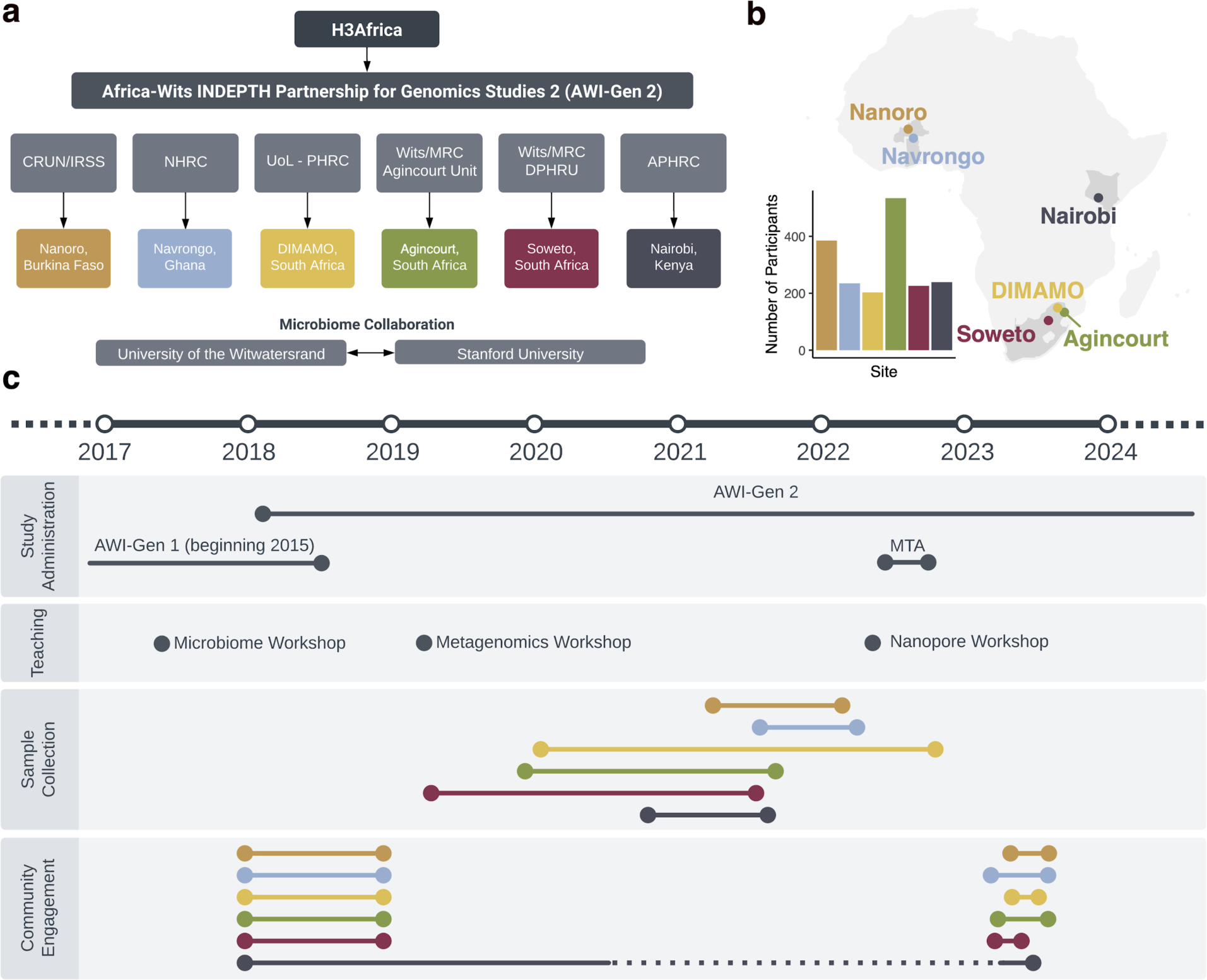
Overview of the AWI-Gen 2 Microbiome study. a) Organizational chart of the AWI-Gen 2 project. The partnership, funded by the National Institutes of Health under the umbrella of the Human Heredity and Health in Africa consortium (H3Africa), includes five Health and Demographic Surveillance Sites (HDSSs) and the Soweto Developmental Pathways for Health Research Unit (DPHRU). The HDSSs and DPHRU are managed by the Clinical Research Unit of Nanoro Institut de Recherche en Sciences de la Santé (CRUN/IRSS), Navrongo Health Research Centre (NHRC), University of Limpopo Population Health Research Centre (UoL - PHRC), University of the Witwatersrand and the South African Medical Research Council (Wits/MRC), and African Population Health and Research Center (APHRC). Researchers from Stanford University and the University of the Witwatersrand lead the microbiome analysis. b) Study site locations and number of participants recruited from each site. c) Timeline of the AWI-Gen 2 microbiome study research activities, including study administration, sample collection, and community engagement. During both AWI-Gen phases, researchers led microbiome and bioinformatic workshops for local researchers. Community engagement preceded sample collection at all sites, and participants with concerning health-related results were referred to their local healthcare facilities in accordance with site-specific protocols. Community engagement in Nairobi continued intermittently throughout sample collection to accommodate roadblocks during the COVID-19 pandemic.

Here, we present the first results from the second phase of the AWI-Gen Microbiome Project. Our pilot studies during the first phase of AWI-Gen established study feasibility, identified preliminary associations between the microbiome and epidemiological transition, and found substantial microbiome novelty^31,32^. In this phase, we aim to evaluate the intersection between geographical, environmental, clinical, and gut microbiome factors in health and disease. First, we randomly sampled 1,820 older adults from well-characterised HDSS populations in six research centres spanning four countries in the west, east and south of Africa (Burkina Faso, Ghana, Kenya, and South Africa). These study centres have widely different population densities, income levels, and disease profiles, with household subsistence strategies ranging from predominantly horticultural subsistence farming to post-industrial. Leveraging extensive clinical and demographic data that were also collected, we identify site or geography as having the strongest effect on microbiome variation and identify microbial taxa with different prevalence patterns across the six sites. We assemble thousands of prokaryotic and phage genomes, including hundreds for *Treponema succinifaciens,* a hallmark bacterial species previously described as absent in industrial populations. Finally, we consider infectious diseases that are relevant to the study communities, and find HIV-associated differences in microbiome composition that are consistent across various sites, but which differ from HIV-associated differences described in studies from HICs. Altogether, this study demonstrates the importance of investigating the gut microbiome in undersampled populations, provides a framework for equitable microbiome research, and represents the largest population-representative profile of African gut microbiomes to date.

## Results

### Study Cohort

Single stool samples were collected from each of 1,820 participants between the ages of 42 and 86 years from rural villages in Nanoro, Burkina Faso^33^ (*n*=382), Navrongo, Ghana^34^ (*n*=235), the Agincourt-Bushbuckridge subdistrict in South Africa^35^ (*n*=533), and Dikgale, South Africa^36^, in which the HDSS is now called DIMAMO (*n*=203), from the township of Soweto, South Africa (*n*=226), and from the Korogocho and Viwandani urban informal settlements in Nairobi, Kenya^37^ (*n*=237). Participants were selected as a cross-sectional representation of the older adults in populations in the catchment areas of each HDSS (see Supplementary Methods). The six study communities span rural, peri-urban, and urban areas, and therefore have in some cases drastic differences in population density, water sanitation, access to healthcare, and disease profiles (**Table 1**). Briefly, the Nanoro and Navrongo study centres are in rural regions of western Africa where subsistence farming and cattle-keeping are dominant subsistence strategies. The Agincourt and DIMAMO study centres both largely consist of semi-rural villages in regions of South Africa, which are undergoing rapid epidemiological transition. Soweto is a district within the city of Johannesburg, which under apartheid was designated as an area for black people to live; employment in Soweto is often related to business, retail, and industry, but unemployment among women remains above 60% ^38^. The Nairobi study centre samples from two urban informal settlements in Nairobi, Kenya, where population density is very high and residents have limited access to piped water and sanitation.

**Table 1.** Study centre characteristics.

|  | Nanoro,<br>Burkina Faso | Navrongo,<br>Ghana | DIMAMO,<br>South Africa | Agincourt,<br>South Africa | Soweto,<br>South Africa* | Nairobi, Kenya |
| --- | --- | --- | --- | --- | --- | --- |
| **HDSS Catchment Area, km2** | 594 | 1,675 | 545 | 420 | 200 | 6 |
| **HDSS Catchment Area**<br>**Population, thousands** | 63 | 156 | 36 | 115 | 1,200 | 75 |
| **Population density per km2** | 105 | 91 | 113 | 274 | 6,357 | 14,833 |
| **Map coordinate** | 12.57N, 1.92W<br>12.72N, 2.31W | 10.89N, 1.09W | 23.65S, 29.65E<br>23.90S 29.85E | 24.50S, 31.08E<br>24.56S 31.25E | 26.24S, 27.84E | 1.25S, 36.89E<br>1.31S 36.87E |
| **Altitude, m** | 313 | 196 | 1,250 | 400-600 | 1,632 | 1,790 |
| **Household size, median (IQR)** | 12 (7, 20) | 6 (5, 7) | 5 (3, 7) | 5 (3, 7) | 4 (2, 6) | 4 (2, 6) |
| **Electricity (%)** | 3.9% | 33.9% | 98.0% | 99.6% | 98.2% | 82.9% |
| **Age, median (IQR)** | 50 (48, 53) | 55 (52, 61) | 53 (50, 57) | 61 (54, 68) | 54 (50, 58) | 50 (48, 53) |
| **Prevalence of HIV** |  |  |  |  |  |  |
| Seronegative, n(%) | - | - | - | 426 (80.2%) | 165 (75.7%) | 216 (90.4%) |
| Seropositive, +ART, n(%) | - | - | - | 60 (11.3%) | 50 (22.9%) | 19 (8.0%) |
| Seropositive, -ART, n(%) | - | - | - | 45 (8.5%) | 3 (1.4%) | 4 (1.7%) |

\* The Soweto study area is managed by the SAMRC/Wits Developmental Pathways to Health Research Unit, not an HDSS
Household size, electricity, age, HIV, and BMI statistics are specific to the AWI-Gen 2 Microbiome Project study populations.
For normally distributed parameters, values represent the mean ± SD; for not normally distributed parameters, values represent the median with IQR (P25, P75). Categorical variables are represented by counts and percentages.
The table above summarizes the catchment area of the sites' surveillance area. Note that the catchment area of the site does not always correspond exactly to administrative units for which census data is available. See the supplementary methods for a full description of the sites and inclusion criteria.

The AWI-Gen Phase 2 study is a population cross-sectional study of adults aged 32-98 (99% between 41 and 84). Pregnant women and people who had been resident for less than 10 years were excluded. At the point of recruitment into AWI-Gen, only one person per household was included. Inclusion criteria for the microbiome study included self-identified female sex, and participation in the larger AWI-Gen Phase 2 study. In total, 43% of women in AWI-Gen 2 were semi-randomly included (see Supplementary Methods). The majority of participants were women (*n* = 1,803), and a small number of men (*n* = 17) were sampled as well. The focus on recruitment of women was motivated by downstream interest in combining these data with a companion AWI-Gen 2 study focused on menopause. Despite limited microbiome differences between sexes (**SFig. 1**), we excluded samples from men for all site comparisons presented below; however, given the poor representation of these populations in existing microbiome studies, samples from men were included in the genome catalogues presented below. Participants completed the AWI-Gen 2 questionnaire with guidance from a supervised field worker, and donated blood, urine, and stool samples. For gut microbiome analysis, DNA was extracted from each stool sample, followed by 2×150 base pair paired-end sequencing. We generated a median of 44.16M (range 27.48M to 104.79M) reads per sample, with a median of 31.20M (range 17.80M to 72.95M) reads remaining after quality control and removal of human reads (see Methods). For an extended description of each study centre and recruitment methodology, see the Supplementary Methods.

### Gut microbiome composition of AWI-Gen 2 participants

We first characterized the overall taxonomic composition and microbial diversity in the study populations. We performed taxonomic classification with mOTUs3^39^, using the GTDB-based taxonomy^40^. After clustering samples by overall microbiome composition, we observe the primary axis of variation to capture a trade-off in relative abundance between the phyla Bacteroidota and Firmicutes A (**Figure 2ab, SFig. 2**), and to correlate with the abundance of the archaeal phylum Methanobacteriota. The second principal coordinate captured study site differences (PCo2; **Figure 2a**), generally ordering the samples along a gradient corresponding to site population densities, subsistence strategies, environments, and sociodemographic factors. The exception to this gradient is Nairobi, Kenya, which is a dense urban site, yet falls in the middle of the gradient. This second axis is correlated with the abundance of the phyla Spirochaetota and Elusimicrobiota, phyla that have been described to decrease in relative abundance with industrialization^41^ (**Figure 2b**), as well as with the observed prokaryotic richness.

**Figure 2.**
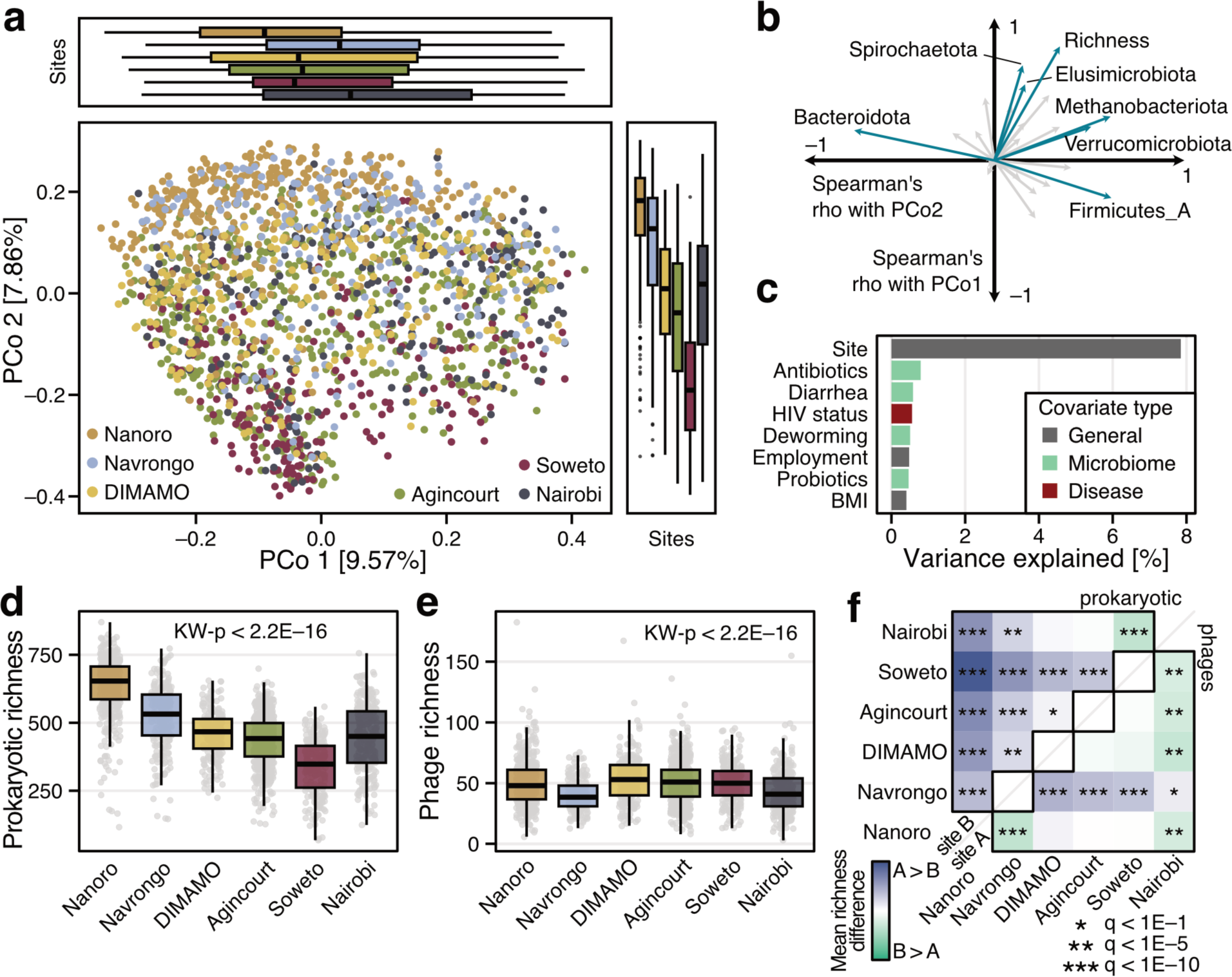
Microbial composition and diversity in the AWI-Gen 2 Microbiome cohort. **a**) Principal coordinate analysis of all samples based on Bray-Curtis distance on species-level prokaryotic profiles. Study site is colour-coded and the boxplots on the side and above show the samples per site projected onto the first and second principal coordinate. **b**) The Spearman correlation coefficient (Spearman’s rho) between principal coordinate values and relative abundance of prokaryotic phyla (and prokaryotic richness) is indicated by arrows. Phyla with an absolute correlation coefficient higher than 0.4 for either of the two principal coordinates are highlighted in blue. **c**) The amount of variance in the prokaryotic composition that is explained by various covariates in distance-based redundancy analysis is shown for all covariates that explain more than 0.5% of variance (**SFig. 3**). **d**) Prokaryotic richness (number of prokaryotic species present at ≥1e-04% abundance after rarefaction) per site(Kruskal-Wallis test *p* < 2e-16, n=1796). **e**) Phage richness (number of phage species clusters present in each sample, see Methods) per site (Kruskal-Wallis test p < 2e-16, n=1796). **f**) Pairwise comparisons between sites in prokaryotic and phage richness. Each tile corresponds to the results of a linear model comparing the richness between two different sites. The fill colour indicates if the richness is higher in site A (see x-axis) compared to site B (see y-axis of the heatmap). Stars indicate the significance (tested by ANOVA) after correction for multiple hypothesis testing using the Benjamini-Hochberg procedure (see Methods). Above the diagonal, prokaryotic richness is compared, whereas comparisons in phage richness are shown below the diagonal. For all boxplots, boxes denote the interquartile range (IQR) with the median as a thick black line and the whiskers extending up to the most extreme points within 1.5-fold IQR.

Seeking to identify the geography, disease, and lifestyle factors that have the greatest effect on compositional variation, we performed distance-based redundancy analysis with the available covariates, excluding highly correlated variables (**SFig. 3**, see Methods). As expected, study site explains, by far, the greatest amount of compositional variation (7.85%), followed by other variables inherently related to the microbiome, such as recency of antibiotic use (0.79%), recency of diarrhoea (0.59%), use of deworming medication (0.51%), or probiotics use (0.47%). Interestingly, the only disease-related variable explaining a sizable amount of composition variation is HIV status (0.52%) (**Figure 2c**), illustrating the potential of this cohort to investigate HIV-microbiome associations. Other disease variables included arthritis, obesity, hypertension, and others (**SFig. 3**).

To delve deeper into differences between study sites, we investigated microbial diversity across sites, focusing on prokaryotes as well as phages. Prokaryotic richness was measured as the number of species present in classification data after rarefaction, and phage richness was measured as the number of phages annotated in assemblies from each sample (Methods). Prokaryotic alpha diversity is significantly different between sites (Kruskal Wallis *P* ≤ 2E-16; **Figure 2d**), mirroring the site gradient observed for the second principal coordinate, with Nairobi again falling out of sequence. However, bacteriophage richness does not follow this population density gradient, even though prokaryotic and phage richness generally correlate well (**Figure 2e**, **SFig. 4**). Interestingly, phage richness seems to be fairly similar across sites, with Navrongo and Nairobi showing significantly lower phage richness than the other sites (**Figure 2f**). These findings were independent of sequencing depth and generally similar with reference-based phage profiling method^42^ (**SFig. 4**), hinting at unexplored factors for phage ecology.

Altogether, study site is the dominant factor explaining overall microbiome composition. Microbiome composition and diversity follow a gradient across study sites, ordered roughly by site differences in socio-demographic, lifestyle, and environmental factors, although Nairobi is often an outlier. Disease and medication variables explain smaller amounts of microbiome composition, and HIV status represents one of the largest subsequent contributors to microbiome variation after study site. These findings support the hypothesis that study sites represent far more than geography, including varied subsistence strategies, industrialization levels, healthcare access, and overall adversity that together affect microbiome composition.

### Hallmark taxa differ across study sites

Given the strong effect of study setting on overall microbiome composition, we next identified the prokaryotic taxa that have distinct prevalence and relative abundance patterns between study sites. We clustered prokaryotic species by population prevalence, defining presence in an individual as relative abundance ≥ 1e-04, in each study site and observe several distinct prevalence patterns (**Figure 3a, STable 2**). Some species are ubiquitous across the study sites and appear to be core members of the gut microbiome in these populations regardless of site, such as *Faecalibacterium* or *Blautia* species, while others have prevalence patterns that are positively or negatively correlated with the study site gradient observed previously. As expected, sites with similar subsistence strategies and populations have highly correlated species prevalence (e.g. Navrongo and Nanoro, 0.91; Agincourt and DIMAMO, 0.96). Two interesting exceptions are Nairobi and Soweto, which are both industrial or post-industrial urban sites but have a prevalence correlation of only 0.65, and Nanoro and Agincourt, which are both sites with lower population density, but have different subsistence strategies and a prevalence correlation of only 0.66. Critically, these poor correlations illustrate that ‘urbanization’ or ‘industrialization’, which are commonly cited variables that impact the microbiome in prior research including our own, cannot adequately capture the diverse lifestyle and environmental factors that exist between different urban areas and between different rural areas.

**Figure 3.**
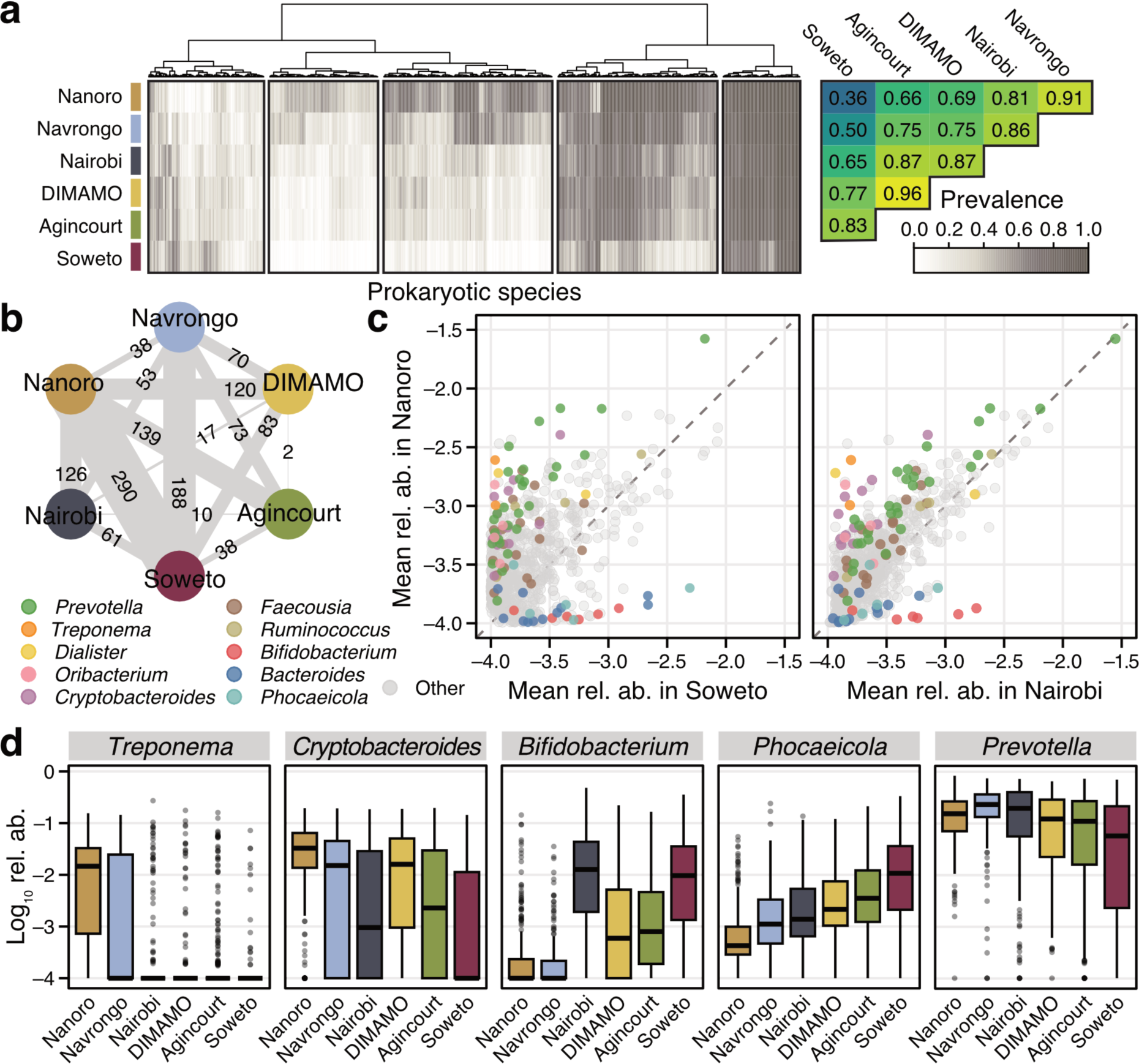
Site comparison reveals patterns of lifestyle-related microbiome transition. **a**) The prevalence per site is shown for all prokaryotic species with prevalence higher than 5% in at least 2 sites (n=886 species), clustered using the Ward algorithm. Spearman correlation between sites is shown on the right. **b**) For the same species as in a), the number of bacterial species that have a high generalized fold change between sites (see Methods) is indicated by the thickness of the edge connecting two sites. **c**) The mean log10-transformed abundance of the same prokaryotic species as in a). Species that belong to the 10 genera with the highest variance in fold change across all sites (see Methods) are highlighted by colours. **d**) The log10 relative abundance of select genera are shown across the six different study sites (sites ordered by the clustering in a). See Fig. 2 for boxplot definitions.

Intrigued by the differences between urban settings and the differences between rural settings, we next identified how many and which prokaryotic species differed most in abundance between sites. As expected, this analysis revealed a pattern similar to the prokaryotic prevalence correlations (**Figure 3b**). We observe very few differentially abundant taxa between sites with similar characteristics (2 species differ between Agincourt and DIMAMO) and several differentially abundant taxa between sites with distinct characteristics (Soweto and Nanoro, 290 species) (**Figure 3b**). We first compared differentially present taxa between Nanoro and Soweto, which fall at two extremes of the site gradient. Species from the genera *Treponema*, *Dialister*, *Prevotella*, and *Cryptobacteroides* are lower in Soweto, whereas species from the genera *Bacteroides*, *Phoecaeicola*, and *Bifidobacterium* are lower in Nanoro (**Figure 3c**). Interestingly, when comparing Nanoro to Nairobi, which like Soweto is highly urbanized and densely populated, *Bifidobacterium*, *Treponema*, and some *Cryptobacteroides* species show similarly extreme differences between the sites, whereas *Phocaeicola* and *Bacteroides* abundance differences are more moderate. The overall relative abundances of these taxa in Nairobi are more comparable to those observed in DIMAMO and Agincourt than to those observed in Soweto (**Figure 3cd; SFig. 5**).

These findings suggest a model for the dynamics of microbiome transition. We observe some taxa such as *Treponema* are easily lost in sites practicing large-scale agriculture or industry, whereas we observe a gradual expansion in abundance of *Bacteroides* and *Phocaeicola* species and a gradual decrease in abundance of *Prevotella* species. Despite numerous reports in other cohorts^43,44^, we do not observe *Prevotella* and *Bacteroides* to be mutually exclusive (**SFig. 5**). We observe interesting taxonomic profiles in participants from Nairobi, many of whom likely migrated to Nairobi from rural parts of Kenya^37,45^ into the informal settlements. The microbiomes of individuals from Nairobi are more compositionally similar to those from participants in DIMAMO and Agincourt (**Figure 3b**) compared to those in Soweto. Furthermore, the microbiomes of individuals from Nairobi have similar abundances of *Prevotella* species as participants in Nanoro, suggesting that participants from Nairobi may have retained a microbial signature from their former rural residences. By contrast, abundances of *Bifidobacterium* species are similarly high and abundances of *Cryptobacteroides* species are similarly low in participants from Nairobi and Soweto, suggesting that these taxa are likely strongly influenced by changing environments.

### AWI-Gen 2 microbiomes contain substantial novelty

African gut microbes are underrepresented in public reference collections such as the Unified Human Gastrointestinal Genome catalogue, and when present, are often sourced from relatively isolated populations with lifestyle practices (e.g. hunting and gathering) that are not representative of the African continent. To identify novel microbial taxa in the AWI-Gen 2 sample collection, we performed metagenomic assembly and binned assembled contigs into metagenome-assembled genomes (MAGs), yielding a total of 69,539 genomes. To condense redundant genomes and build a species set, we took all MAGs with a minimum genome completeness of 50% and maximum genome contamination of 5%, and dereplicated the genomes at 95% average nucleotide identity (ANI). The resulting catalogue of 2,613 MAGs span 19 bacterial phyla (**Figure 4a**).

**Figure 4.**
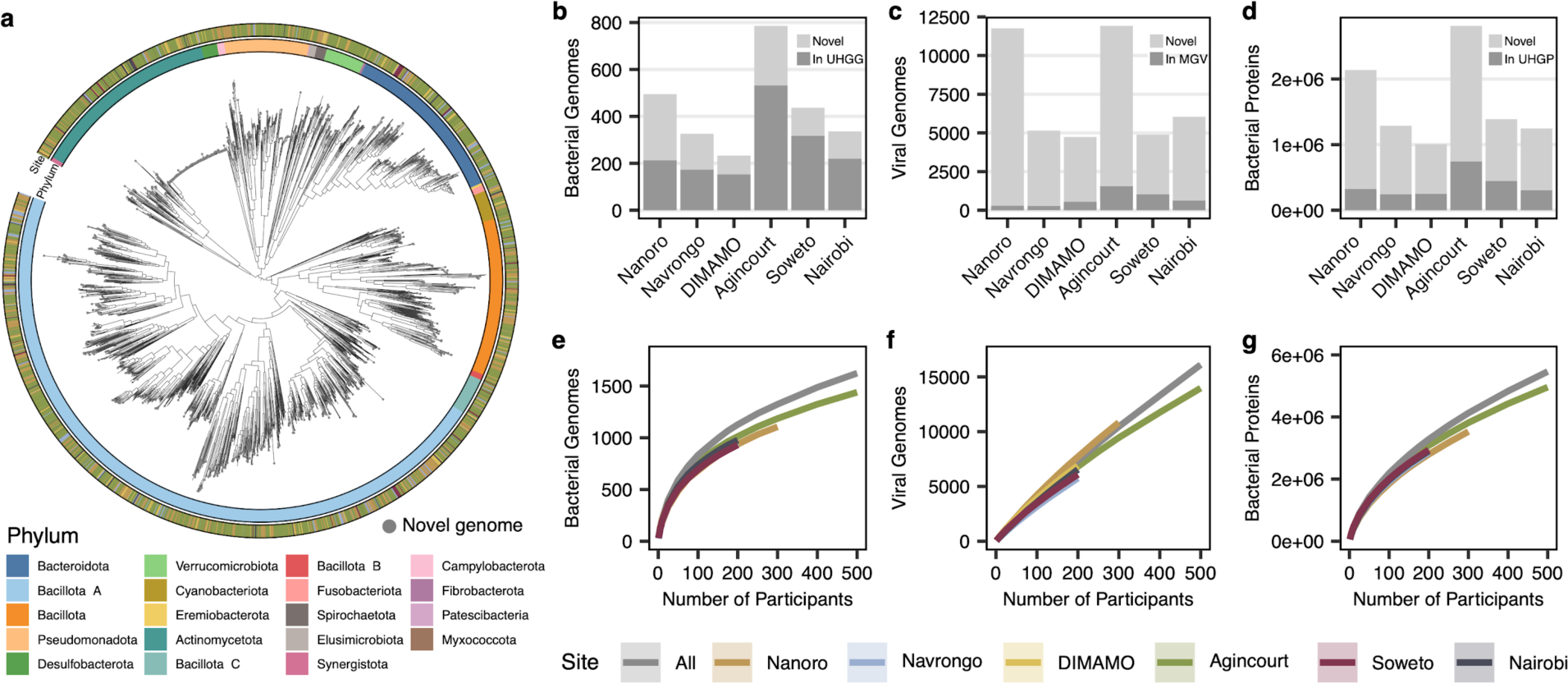
Catalogue of novel microbial features. **a)** Phylogenetic tree of 2,584 de-replicated bacterial metagenome-assembled genomes generated in this study. Outer ring indicates study site of origin, inner ring indicates assigned GTDB phylum, and leaf points indicate genomes that are novel relative to UHGG. Total number of novel and existing **b)** prokaryotic genomes, **c)** viral genomes, and **d)** prokaryotic proteins in the AWI-Gen assemblies, relative to existing databases for each feature. Only representative genomes and proteins after feature clustering are represented. Rarefaction curves of the number of **e)** prokaryotic genomes, **f)** viral genomes, and **g)** prokaryotic proteins detected as a function of the number of individuals sampled, by study site or from the full AWI-Gen sample set (grey). Each random subset was repeated a hundred times, and lines represent the mean feature count and standard deviation.

Although most gut microbiome research focuses on prokaryotic composition and prokaryotic genomes, this sample collection also represents a likely source of viral and functional diversity. Therefore, along with our catalogue of prokaryotic MAGs, we generated a viral genome catalogue from all assemblies clustered at 95% ANI and a protein catalogue from all medium- and high-quality MAGs clustered at 95% amino acid identity. We compared our prokaryotic, viral, and protein catalogues to the current field-standard reference collections: the Unified Human Gastrointestinal Genome (UHGG) catalogue of 4,744 prokaryotic species representatives, the Metagenomic Gut Virus catalogue of 54,118 viral operational taxonomic units, and the Unified Human Gastrointestinal Protein 95 (UHGP95) catalogue of 20.5 million proteins. These comparisons revealed that the AWI-Gen 2 dataset is enriched for novel features, including 1,005 new prokaryotic MAGs relative to UHGG (**Figure 4b**), 40,135 new viruses relative to MGV (**Figure 4c**), and 7.6 million new proteins relative to UHGP95 (**Figure 4d**). Notably, we observe that a greater portion of proteins and viruses are new to reference collections, compared to prokaryotic genomes which are moderately well-represented. Most individual samples yielded several novel prokaryotic and viral genomes and tens of thousands of novel proteins relative to reference collections (**SFig. 6**).

Given the substantial addition of new prokaryotic genomes, viral genomes, and proteins in the AWI-Gen 2 microbiome project, we next explored whether this sample collection has saturated possible metagenomic discovery from these populations. We performed a rarefaction analysis, counting the total number of prokaryotic genomes (**Figure 4e**), viral genomes (**Figure 4f**), or proteins (**Figure 4g**) generated from samples randomly selected from within or across each of the six study sites. We find that no feature has reached saturation and that the addition of new samples continues to yield additional features, especially for proteins and viral genomes. Critically, these results imply that additional measurement of gut microbiomes in these communities and in other African populations will continue to reveal new microbiome diversity.

### Leveraging MAG database to understand bacterial biology

The extensive AWI-Gen 2 microbial genome catalogue enables us to better understand organismal biology, including the biology of organisms that have yet to be extensively studied using standard microbiological techniques. One taxon of particular interest is *Treponema succinifaciens*, a commensal anaerobic bacterial species in the phylum Spirochaetes that is non-spore forming, consumes a wide range of sugars, and produces short-chain fatty acids and succinate^46^. *T. succinifaciens* is thought to be present in rural hunter-gatherer, pastoralist, and agriculturalist populations, and lost in urban populations^16,41^. Previous work in the first phase of AWI-Gen identified that *T. succinifaciens* is indeed present in urban populations^32^; however, we do observe that *T. succinifaciens* prevalence is inversely correlated with population density (**Figure 3d**). Despite the emerging interest in this gut commensal, *T. succinifaciens* dispersal and acquisition is poorly understood.

*Treponema* are difficult microbes to culture^47^, making metagenome-assembled genome catalogues an invaluable resource for better understanding their biology. Only one complete genome from a cultured *T. succinifaciens* is available^46^, and this type strain was isolated from the swine gut^48^. Only 99 *T. succinifaciens* MAGs exist in UHGG, predominantly assembled from human gut samples from populations in Madagascar, Peru, and Fiji. Our genome catalogue includes 249 metagenome-assembled genomes for *T. succinifaciens*, with most genomes originating from sites with low population density (**Figure 5a**). Regardless of site of origin, genome lengths are relatively small (2.51 ± 0.18 megabases; **Figure 5b**), and their shared core genome of 1,578 genes constitute the majority of each genome (69.17% ± 3.93%; **Figure 5c**). Based on the knowledge that *T. succinifaciens* is not spore forming and decreases in prevalence with industrialization, along with the observation that *T. succinifaciens* has a small and relatively compact genome, we hypothesized that *T. succinifaciens* is a host-adapted bacterial species with limited environmental dispersal. If that is the case, we would expect that most individuals acquire *T. succinifaciens* locally, and that *T. succinifaciens* would therefore have a strong signature of phylogeography. Indeed, by comparing *T. succinifaciens* genomes from this study to those from other studies^49,50^, we observe that *T. succinifaciens* genomes cluster by continent or country of origin (**Figure 5d**), a pattern which is also visible within our catalogue alone (**SFig. 7**). This phylogeographic signature supports a hypothesis of local dispersal and acquisition of *T. succinifaciens*, and provides one example of how metagenome-assembled genome catalogues built from populations in understudied areas can be used to investigate bacterial biology.

**Figure 5.**
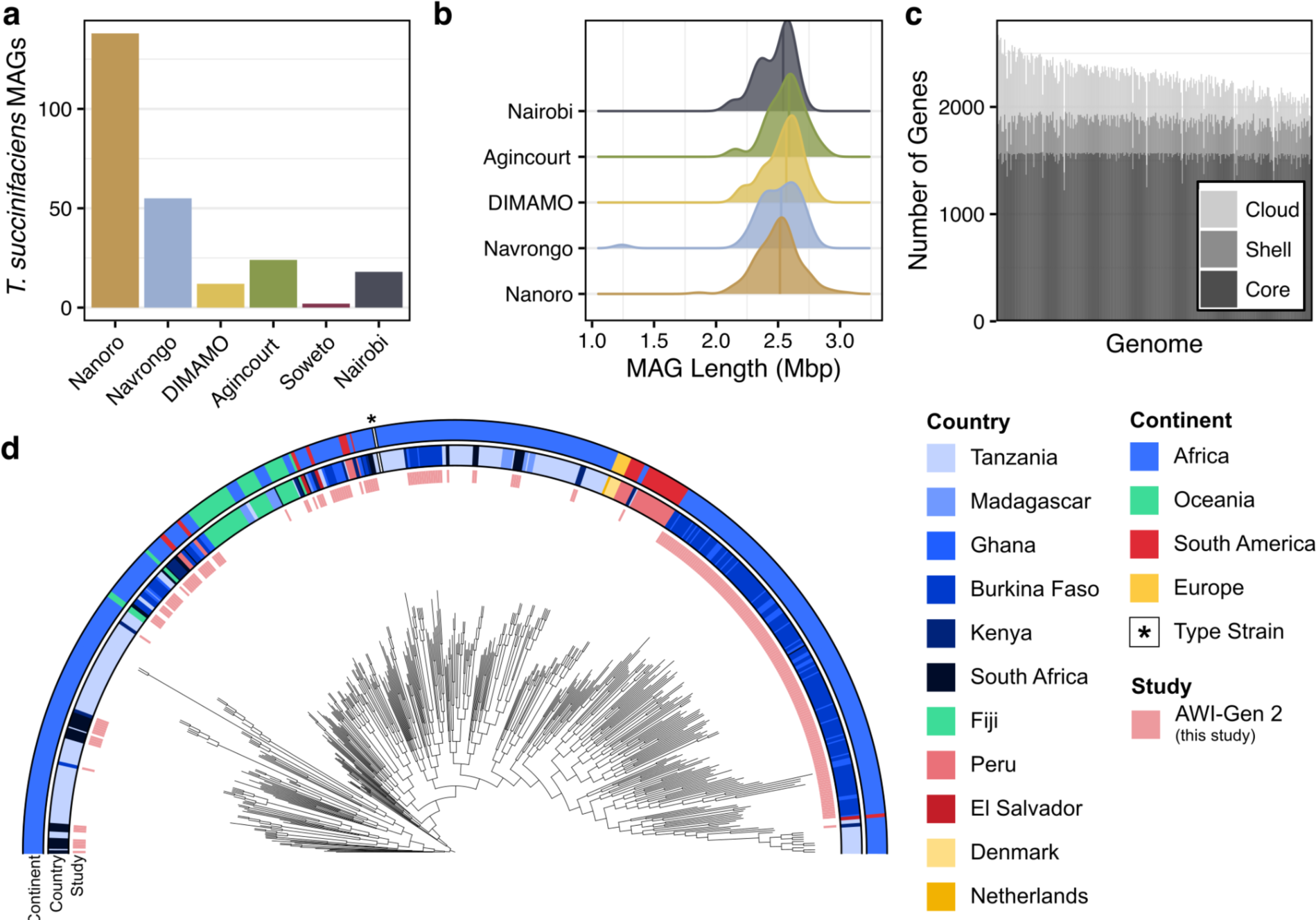
Features of *Treponema succinifaciens* metagenome-assembled genomes (MAGs). a) Number of *T. succinifaciens* metagenome-assembled genomes by study site. b) Distribution of the length, in megabase pairs (Mbp), of each *T. succinifaciens* MAG. MAGs from Soweto are not pictured, as Soweto samples only contained two MAGs. c) Number of genes in each MAG that were classified as core (≥ 80% prevalence), shell (25 ≤ prevalence < 80%), or cloud genes (< 25% prevalence) in the complete MAG set. d) Midpoint-rooted phylogenetic tree of *T. succinifaciens* MAGs from this study (noted in pink inner ring) and public data sets (n = 513 total genomes). Middle ring indicates the country of origin, and outer ring indicates the continent of origin. White line and asterisk indicate the *T. succinifaciens* DSM 2489 type strain reference genome.

### Gut microbiome associations with HIV status

Finally, we investigated the relationship between the gut microbiome and HIV status, as HIV represents one of the biggest public health concerns in the Kenyan and South African AWI-Gen study populations: HIV prevalence estimates in 2022 among individuals aged 15 - 49 is 17.0% in South Africa^51^ and 3.7% in Kenya^52^. Despite advances in antiretroviral therapy (ART), viral suppression is not sufficient to control HIV-related mortality and morbidity^53^, thus motivating deeper investigation into the gut microbiome as a possible mediator. Gut microbiota and their metabolites have been implicated in HIV-related inflammation and immune activation: gut-associated lymphoid tissue serves as a major reservoir for HIV^54,55^, and gut microbial metabolites can promote HIV transcription^56,57^. In turn, HIV infection can diminish epithelial barrier integrity^58^, allowing for microbial translocation that promotes immune activation and chronic inflammation^59,60^. Moreover, obesity has become a major problem for individuals on the latest generation of ART such as dolutegrevir^61^. Very few studies have measured associations between the gut microbiome and HIV status in African populations^13,62–64^, where baseline microbiome composition and disease profiles are distinct from those observed in HICs.

The AWI-Gen 2 project represents a unique opportunity to understand the associations between the microbiome and HIV in a population-representative sampling. We compared microbiome composition between women living with HIV (people living with HIV; PLWH) and seronegative (HIV−) women in Agincourt (PLWH *n*=60; HIV− *n*=342), Soweto (PLWH *n*=50; HIV− *n*=165), and Nairobi (PLWH *n*=19; HIV− *n*=214) (**Figure 6a**). HIV status was not assessed in Nanoro and Navrongo because of low population prevalence in those populations, and DIMAMO was excluded for this comparative analysis, since only six individuals were found to be living with HIV. Most PLWH are undergoing ART as standard therapy to manage their HIV infection. Our dataset also included participants with positive HIV status, but not self-reporting as receiving ART, possibly because they learned of their HIV diagnosis in the course of participation within AWI-Gen 2. Due to the low number of ART-naive PLWH in our dataset, we focused our analysis only on HIV- and ART+ PLWH participants (see **SFig. 8** for comparison between ART+ and ART− PLWH).

**Figure 6.**
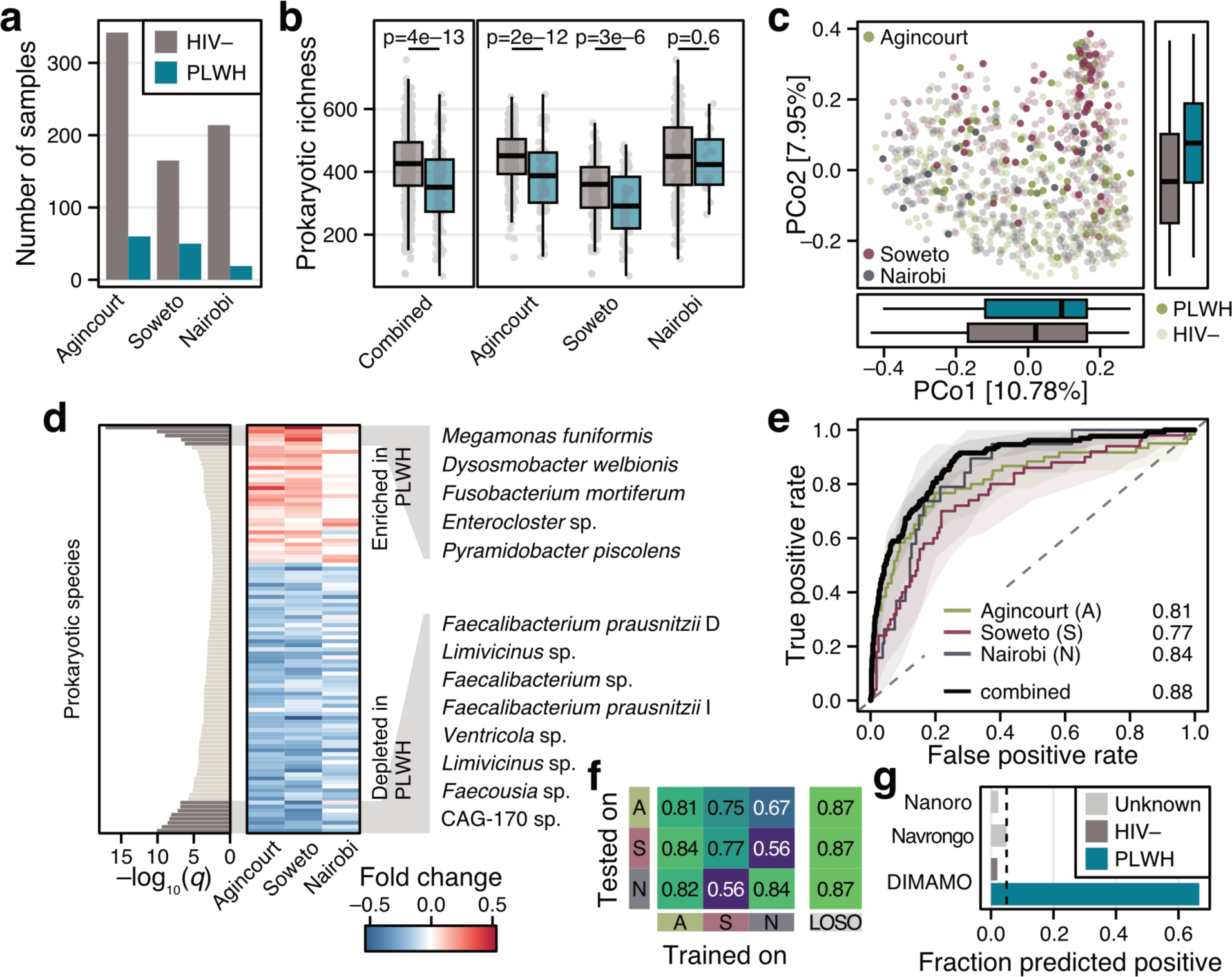
Microbial composition and diversity in people living with HIV. **a)** Number of seronegative individuals (HIV-) and people living with HIV (PLWH) on antiretroviral treatment included for analysis. **b)** Prokaryotic richness (number of prokaryotic species present at ≥1e-04% abundance) by site and HIV status. Points represent individual samples. Differences in alpha diversity for each individual site were tested with ANOVA and for all sites combined with a linear mixed effect model accounting for site as a random effect. **c)** Principal coordinate analysis of all samples based on Bray-Curtis distance on species-level prokaryotic profiles. Points represent individual samples, coloured by study site, and PLWH are shaded. Boxplots on the side and below show the samples by HIV status projected onto the first and second principal coordinate. **d)** Differentially abundant species as determined by a linear mixed effect model accounting for confounders (see Methods). Species with q-value < 0.01 are shown and species with q-value < 1e-05 are annotated (see **STable 3** for the full list). Shading indicates site-specific abundance fold change between seronegative individuals and PLWH. **e)** Receiver-operating characteristic (ROC) for machine learning models trained to distinguish HIV status on samples from each site or for all data combined. Shaded areas indicate the 95% confidence interval and numbers indicate area under the ROC curve (AU-ROC). **f)** AU-ROC values for machine learning model evaluation. Models trained on participants from each site were applied to the data from other sites and the external predictions were evaluated via AU-ROC (see Methods). In the leave-one-site-out (LOSO) validation, models were trained on data from two sites and validated on the left-out site (e.g. model trained on data from Soweto and Nairobi was evaluated on data from Agincourt). **g)** Fraction of samples from other sites predicted to be positive calibrated at a 5% false positive rate (indicated by dashed black line, see Methods). For DIMAMO, HIV status is known and therefore, the false positive rate and the true positive rate can be evaluated. Note that serostatus is not known for individuals in Nanoro and Navrongo but is expected to be below 2%. See Fig. 2 for boxplot definitions.

To first compare the microbiome of PLWH and HIV− individuals on an overall composition level, we compared alpha and beta diversity metrics. Consistent with previous descriptions^13,65^, alpha diversity is lower in PLWH overall, and this difference is also detected within each study site (**Figure 6b**). In terms of the beta diversity, HIV status significantly varies over both principal coordinate axes (PCo1 *P*=0.05, PCo2 *P*=1.5e-11, Wilcoxon test) but is again outweighed by site differences (PCo1 *P*=6.97e-11, PCo2 *P*=1.12e-26, Kruskal Wallis test; **Figure 6c**).

To identify specific taxa that vary with HIV status, we performed differential abundance testing with a linear mixed effect model that allowed us to account for confounders such as site, antibiotic treatment, or recency of diarrhoea. After correcting for multiple testing, 101 prokaryotic species showed significant differences with HIV status (q-value < 0.01, see **Figure 6d** and **STable 3**), the majority of which seem to be lost in PLWH, agreeing with the finding of lower prokaryotic richness in PLWH. Overall, the effect sizes generally agree across study sites, with Nairobi exhibiting smaller effect sizes due to a lower number of PLWH. Some of the most significantly associated (q-value < 1e-05, **Figure 6d**) taxa have been associated with HIV status in other cohorts: *Faecalibacterium prausnitzii* is a known anti-inflammatory species and has been negatively associated with HIV status previously^66^, and the genus *Fusobacterium* has well-described pro-inflammatory associations with HIV and other diseases^67,68^. Interestingly, other taxa that are negatively associated with HIV, including *Ventricola sp.*, *Faecousia sp.*, and *Limivicinus sp.* are poorly described and better represented in metagenomic studies focused on livestock^69^. Taxa that are positively associated with HIV, including *Dysosmobacter welbionis* and *Megamonas funiformis*, have conflicting associations with inflammatory disorders^70,71^ and remain poorly characterized. These results highlight the value of investigating microbiome and disease associations in broader cohorts, as many of these highly associated taxa are not present in studies conducted in HICs.

To explore whether microbiome composition could be used to predict HIV status, we trained machine learning models for each site individually and for all data combined (see Methods, **Figure 6ef**). These models achieved accurate distinction between HIV- and PLWH individuals in all sites, yielding area under the receiver-operating-characteristics (AU-ROC) > 0.75 in all cases. When transferred across sites, only the model trained on data from Agincourt maintains high classification accuracy (see **Figure 6f**), which is probably the result of a larger data set available for model training. In line with this hypothesis, when data from two sites were combined in a leave-one-site-out (LOSO) validation approach, samples from the left-out site are still accurately classified (AU-ROC = 0.87 in all cases), even when samples from Agincourt are not used for training at all. As another test for generalization, we calibrated the model trained on all data to an internal 5% false positive rate (FPR, see Methods) and applied it to the samples from Nanoro, Navrongo, and DIMAMO (**Figure 6g**). Even though information about HIV status in Nanoro and Navrongo are not available, we can still assess the fraction of samples predicted to be HIV positive, which we would expect would not exceed 5% given the population HIV prevalence in these sites and our model calibration. Indeed, the model predicts very few samples to be HIV positive (2.4% in Nanoro, 5.1% in Navrongo), highlighting its specificity, and correctly classifies two thirds of the HIV-positive samples from DIMAMO (**Figure 6g**).

Overall, we find HIV-associated microbiome differences to be consistent across study settings, despite the strong effect of study context on overall microbiome composition. Some strongly associated taxa have been described in the context of HIV previously, but we also identify taxa that have not been well-described in human gut microbiomes or in microbial associations with disease.

## Discussion

In 2007, the Human Microbiome Project set the goal of measuring the human microbiome and understanding its contribution to disease^72^. Subsequent studies have built upon this goal, studying the relationship between the human microbiome and disease in large cohorts in HIC such as Israel^73^, the Netherlands^5^, and Finland^6^. Here, seventeen years later, the AWI-Gen 2 Microbiome Project is a landmark research effort that extends these goals to additional diverse LMIC populations in Africa. This project represents the largest cross-sectional and population-based survey of microbiome composition in relation to human health, environment, and disease in low- and middle-income settings, and will prove invaluable in future microbiome discovery research.

We present the AWI-Gen 2 Microbiome Project, a collaborative research effort that has measured the gut microbiome in a cross-sectional population sample from six African communities across four countries. The AWI-Gen 2 study centres encompass a wide range of environments, including rural and predominantly horticultural areas (Nanoro, Burkina Faso and Navrongo, Ghana), rapidly transitioning rural areas (Agincourt and DIMAMO, South Africa), urban industrial informal settlements (Viwandani and Korogocho settlements in Nairobi, Kenya), and an urban post-industrial settlement (Soweto, South Africa). Paired with extensive participant data, which include human genotype data, blood and urine biomarkers, disease risk outcomes, and demographic and environmental information, the AWI-Gen 2 Microbiome Project represents the largest research effort of its kind in LMIC settings and fills a critical gap in measuring the relationship between the gut microbiome and diverse health, geographic, and environmental variables. This unique study design has supported the discovery of new findings that are unattainable in other datasets.

First, the diversity of study settings enables comparison between populations that span a wide range of subsistence strategies and resource access. We find that study context has a strong effect on microbiome variation, with alpha diversity and prevalence of several taxa correlating with gradients in population density and resource access. Unexpectedly, we observe differences between sites that have similar subsistence strategies and industrialization levels. For example, some taxa (*Bifidobacterium*, *Cryptobacteroides*) found in microbiomes from Nairobi have abundance levels similar to abundances in Soweto, whereas other taxa (*Prevotella*, *Bacteroides*, *Phoecaeicola*) have abundances that are more typical of the four rural and semi-rural sites. Data from the HDSSs can help us contextualize these findings: the Korogocho and Viwandani informal settlements in Nairobi are characterized by high in- and out-migration rates, with approximately a quarter of the population migrating into or out of the settlements each year^37,45^. Approximately 60% of residents migrated from rural areas, and 50-80% of residents revisit their rural homes semi-regularly^45^. The participants in AWI-Gen 2 have all been residents of their respective catchment areas for at least ten years; however, the taxonomic signatures in microbiomes from Nairobi participants are indicative of microbiomes of communities in transition, highlighting taxa that persist in the gut microbiome throughout urban migration, as well as the taxa that are lost or gained during this transition. The intermediate microbiome compositions of participants from Agincourt and DIMAMO may similarly reflect lifestyle: these study centres capture villages that are in the midst of industrialization, and experience high rates of circular migration to urban centres in South Africa for employment, a legacy of former apartheid policy in South Africa^74^. Microbiome changes with international immigration have been observed previously^21^, but here we identify compositional shifts that likely occur as a result of within-region migration and social and epidemiological transition. Studying these populations is critical, as the vast majority of the world’s population reside in LMICs, and over a billion people globally are estimated to live in urban informal settlements^75^. It is a common oversimplification in many microbiome studies to use industrialization or urbanization as binary variables. Our findings instead motivate the incorporation of population demographics, and illustrate the importance of comparing populations within LMICs.

Next, this large cohort of individuals from understudied regions enabled extensive microbial discovery. The bacterial, viral, and protein novelty found in this study emphasizes the severe underrepresentation of gut microbiome data from LMICs in microbiome catalogues. This MAG collection enables investigation into microbes that are absent from reference databases due to low prevalence in high-income countries. For example, here we identify that *T. succinifaciens* has a strong phylogeographic signature, which suggests that this gut commensal is likely acquired locally and has limited geographic dispersal. *T. succinifaciens* is considered a hallmark taxon that is lost in urban individuals^41^. Here and previously^32^, we have found *Treponema* in the guts of urban individuals, perhaps due to high rates of population-level circulatory migration between urban and rural areas. If so, these data may suggest that *T. succinifaciens* is not irreversibly lost in industrial populations, as previously described^76^. This finding illustrates the utility of LMIC population representation in microbial reference collections.

Finally, the population-representative enrolment in AWI-Gen supports the capture of population-level trends and association between microbiome composition and disease. Here, we describe the microbiome signature of HIV infection in Nairobi, Kenya and Agincourt and Soweto, South Africa. HIV prevalence is high in South Africa and Kenya, and improved viral load management with antiretroviral therapy is not sufficient to protect against HIV-associated comorbidities^53^. We find several taxa that are enriched in PLWH receiving ART relative to seronegative participants across sites, including taxa that have not been well-described in the context of HIV and inflammation. It is important to acknowledge that these microbial signatures and enriched taxa may be due to HIV infection itself, or due to medication or other confounding factors that were not measured. These results further support the importance of continued microbiome research in LMICs, as existing disease associations are likely not broadly portable and more research is necessary to disentangle the effects of HIV infection and other confounding variables on microbiome composition.

We emphasize that the AWI-Gen 2 Microbiome Project is not an exhaustive representation of any country or region. As indicated herein, there is tremendous diversity within LMIC populations, and population density alone is not a sufficient indicator of overall microbiome composition. Rather, the microbiome field needs to improve representation of LMIC populations, both to maximize sampling diversity when drawing microbiome associations and to ensure portability of study findings to broader populations. Even within this study, we focus on older adult women and specifically highlight key variables that explain the greatest amount of microbial variation, leaving several disease and lifestyle variables open for future investigation. Although sex differences are generally considered to have less impact on microbiome composition than other factors^77^, many of these studies have been conducted in HICs and therefore do not capture health and lifestyle differences that may exist between sexes in LMICs. Future studies can expand the presented framework to evaluate compositional differences between sexes and across age groups. Further studies can also incorporate microbiome measurements that were not included here. For example, eukaryotic profiling would enable exploration of environmental factors related to pathogenic eukaryote acquisition, or deeper investigation into the complex relationship between eukaryote infection, immune activation, and gut microbiome composition. Total microbial concentration quantification could overcome biases of compositional data to shed light on whether the taxonomic shifts observed in this study are due to blooms or losses of specific taxa, and could be used to identify whether total microbial concentration is related to any health variables measured in this study. Future work can leverage the extensive AWI-Gen 2 participant data and additional methods of microbial quantification to investigate the complex interplay between the microbiome and host genetics, environmental exposures, health status, and participant demographics.

We strove to conduct the AWI-Gen 2 Microbiome Project ethically and equitably, taking into account recommendations for ethical research and equitable partnerships between research groups^24,26,78,79^. The AWI-Gen Collaborative Centre provides an effective example of these principles. First, AWI-Gen works closely within each study community. The study hires field workers locally through the community-embedded HDSS and DPHRU infrastructures, and study staff host community advisory group and community discussions prior to study onset, address community concerns as they arise, and engage upon study completion for feedback of results. Through community discussions and population surveillance, the research team can identify pressing health issues within each study centre and ensure that research questions focus on community needs. The microbiome component of AWI-Gen represents a strong scientific partnership between Stanford University in the USA and University of the Witwatersrand in South Africa. Trainees and faculty from both research groups contributed to study design and data analysis, and a trainee from each institution has participated in a one-year research exchange with joint membership from both institutions. Further, the team has led three training workshops on microbiome and bioinformatics to support strengthening of genomics research capacity in South Africa. Altogether, this study illustrates that equitable research and impactful science do not represent a ‘zero-sum’ tradeoff, but in fact lead to more robust research with benefit-sharing among all stakeholders.

The AWI-Gen 2 Microbiome Project represents a substantial contribution to measuring the composition and diversity of human gut microbiomes from diverse populations around the globe. The data within this study provide extensive opportunities for continued exploration, including identifying microbiome and disease associations, measuring human genetic contributions to microbiome composition, and defining lifestyle factors that shape microbial community assembly. Future studies can leverage these data, along with the foundation for community-engaged and equitable research described herein, to close the gap in global representation in microbiome research. Moreover, there is every reason to anticipate that the platforms established and the findings that emerge will enhance disease management, health and wellbeing among communities living in a diversity of contexts.

## Methods

### Ethics approval

Human subjects research approval was obtained (University of the Witwatersrand Human Research Ethics Committee Clearance Certificate No. M170880, M2210108), and ethics approvals were also obtained at each study centre. Informed consent was obtained from participants for all samples collected. Every participant was provided with an information sheet and consent documents, either in English or translated into the local language. Participants had opportunities to discuss concerns with the interviewer, and participants who could not read or write had documents read aloud with a witness^30^. The Stanford University Institutional Review Board deemed that the de-identified data transferred to Stanford University do not constitute human subjects research and thus did not require an additional ethics approval beyond the human subjects research approval obtained at the University of the Witwatersrand.

### Community engagement

Each study centre conducted pre-study engagement prior to recruitment during both AWI-Gen 1 and AWI-Gen 2, adapting to the local contexts to engage with community members and discuss feedback and concerns related to the study. For example, in DIMAMO, South Africa, pre-study engagement involved meeting with tribal leaders, the community advisory team, and community representatives. In Navrongo, Ghana, the community engagement team visited chiefs and elders of the various study communities and informed them of the proposed study, and followed up with a community sensitization durbar prior to AWI-Gen 1 with a larger audience of chiefs, elders, and people of the study communities. The community durbar was excluded from AWI-Gen 2, due to the ongoing COVID-19 outbreak. In Nairobi, Kenya, the community engagement team held several consultative meetings with members of the community advisory committee, village elders, community health volunteers, and AWI-Gen study participants before, during, and after the study. The village elders and community health volunteers were critical in mobilizing study participants who could not be reached by phone. Questions from participants were related to how blood and stool samples would be used and why the study was focused on women. If during recruitment and sample collection there were notable health concerns (e.g. hypertension), the participants are referred into their clinical healthcare service infrastructures. These mechanisms and processes varied from country to country and for sites within a country, depending on resources and local context.

### Study design and cohort selection

Inclusion criteria included prior participation in the AWI-Gen 1 study^27^ and continued participation in the AWI-Gen 2 study. This AWI-Gen 2 microbiome study is a companion study to an AWI-Gen 2 menopause study, and so only participants self-identifying as female were surveyed for the microbiome sub-study. A small number of men in Nanoro, Burkina Faso were recruited due to a fieldwork mix-up. Given the understudied nature of these populations, we did not fully exclude samples from men in downstream analyses; rather, samples from men were excluded from site comparisons and disease associations, but included when cataloguing genomic novelty. Participants were chosen semi-randomly from the overall AWI-Gen 2 participant pool, with additional measures taken to ensure a cross-section of individuals with respect to menopause status and hypertension. See Supplementary Methods for extended recruitment details.

In Soweto, Nairobi, and Nanoro, participants came to central locations for interviews and biomarker collection. Participants were given stool sample collection kits that were either collected the same day or collected from their homes or at a central location in the following days. At the Navrongo, DIMAMO, and Agincourt study centres, participants were visited in their homes for interviews and biomarker collection. Participants were given a stool sample collection kit to use at their home, which was collected by fieldworkers within 24 hours.

Single stool samples from each participant were self-collected and placed in the OMNIGene GUT OMR-200 Collection Kit by the participants (DNA Genotek). Samples were frozen at study centres and shipped frozen to a central laboratory in Johannesburg, South Africa, where they were thawed, aliquoted into cryovials, and stored at −80°C. After obtaining necessary exportation and importation permits, samples were shipped on dry ice to the United States for downstream processing. Prior analysis was conducted to ensure that storage and shipping conditions would not significantly affect measured microbial composition^80^. Participant metadata, including age, demographic information, health history, and blood biomarkers were collected as part of the larger AWI-Gen 2 project, with methods similar to those used in AWI-Gen 1^30^.

### DNA extraction and metagenomic sequencing

DNA was extracted from samples using the QIAamp PowerFecal Pro DNA Kit (QIAGEN; Cat. No. 51804) from 300 µl of stool sample according to manufacturer’s instructions. Bead beating was performed for ten minutes at 30 Hz, followed by rotation of the adapter and ten additional minutes of bead beating using a TissueLyser II (QIAGEN; Cat. No. 85300) using a 2 ml Tube Holder Set (QIAGEN; Cat. No. 11993), and DNA extractions were eluted with C6 Elution Buffer in a final volume of 80 µl. DNA concentration was quantified by spectrophotometer using the DropSense 96 platform (Trinean; Cat. No. 10100096). Every extraction batch of 96 samples included one water blank as a negative control and one mock community aliquot (Zymo Research; Cat. No. D6300) as a positive control. Samples were evaluated for concentration, integrity, and purity prior to library preparation using the 5400 Fragment Analyzer System (Agilent; Cat. No. M5312AA). Metagenomic libraries were prepared using the NEB Ultra II kit (NEB; Cat. No. E7645L) according to manufacturer’s instructions. Library concentration was quantified using qPCR and fragment length distribution was analysed using a 2100 Bioanalyzer (Agilent; Cat. No. G2939BA). Libraries were pooled and 2×150 bp reads were generated using the NovaSeq 6000 platform (Illumina; Cat. No. 20012850).

### Metagenomic read pre-processing and taxonomy profiling

Metagenomic reads were deduplicated using htstream SuperDeduper v1.3.3 with default parameters, trimmed using TrimGalore v0.6.7 with a minimum quality score of 30 and a minimum read length of 60. Reads aligning to version hg38 of the human genome were removed using BWA v0.7.17^81^. Metagenomic reads were taxonomically profiled using mOTUs v3.0.3^82^ and counts were distributed to GTDB^40^ species using the GTDB_v207 mapping file available as part of the mOTUs database. All samples were used for metagenome assembly and novel feature discovery (n = 1,820). Samples from males and samples with high percentages of human reads (percentage of human reads ≥ 70%, *n*=4 samples) were excluded from classification-based analyses and site comparisons.

### Microbial diversity and site differences

To measure microbial alpha diversity, species counts were rarefied to 5,000 using the `rrarefy` function available through the vegan R package v2.6-4^83^. Alpha diversity was measured as prokaryotic richness (number of species with relative abundance ≥ 1e-04 after rarefaction). Phage richness was measured as the number of unique phages annotated in whole-metagenome assemblies from each sample using VIBRANT v1.2.1^84^ (without rarefaction, see “Metagenome assembly and external dataset comparison” below). Orthogonal phage profiling using read-based classification (see **SFig. 4**) was performed with Phanta v1.1.0^42^ using the combination of MGV and UHGG as the reference database. Differences between alpha diversity metrics across sites were tested with a linear model, using the `anova` function from base R to estimate the significance of the difference.

Beta-diversity was calculated on the Bray-Curtis distance using the `vegdist` function from vegan^83^ and the `pco` function from the labdsv R package v2.1-0^85^. To assess the amount of variance explained by covariates, distance-based redundancy analysis was performed with the `dbrda` function from vegan. In an iterative manner, the covariate explaining the highest amount of variance was added to the model formula. To reduce redundancy of highly correlated covariates, all available covariates were transformed into numerical values (using ordinal factors, whenever applicable) and the Pearson correlation between covariates was calculated. In cases of highly correlated covariates (Pearson’s *r* ≥ 0.8), the covariate that explained the higher amount of variance in the prokaryotic composition was chosen for the iterative model (**SFig. 3**).

The difference between sites for individual taxa was calculated using a generalized fold change^86^. In short, instead of comparing the median (the 50% quantile) between distributions, the generalized fold change is the mean of the differences between two distributions at multiple quantiles and can therefore resolve differences also in low-prevalence taxa. **Figure 3b** shows the number of taxa for which the generalized fold change between sites exceeds the 90% quantile of all pairwise site comparisons across all prokaryotic species.

The number of samples for these analyses (displayed in **Figure 2** and **3**, and associated supplements) was distributed across the different sites as follows: Nanoro, n=382; Navrongo, n=218; DIMAMO, n=201; Agincourt, n=532; Soweto, n=226; Nairobi, n=237.

### Metagenome assembly and external dataset comparison

All samples (n = 1,820), including samples for male participants, were included in metagenomic assembly analyses (Nanoro, n=384; Navrongo, n=235; DIMAMO, n=203; Agincourt, n=533; Soweto, n=226; Nairobi, n=239). Metagenomic reads were assembled using megahit v1.2.9^87^ and assembly quality was assessed using QUAST v5.2.0^88^. Protein-coding genes were predicted from assemblies using prodigal v2.6.3^89^. Metagenomic assemblies were binned into draft genomes using MetaBAT v2.5^90^, CONCOCT v1.1.0^91^, and MaxBin v2.2.7^92^, and subsequently dereplicated and aggregated on a per-sample basis using DAS Tool v1.1.6^93^. Bin quality was assessed using CheckM v1.2.2^94^. To create a dereplicated genome set, metagenome-assembled genomes were dereplicated using dRep v3.4.3^95^, filtering to only include genomes with a minimum CheckM completeness of 50% and maximum CheckM contamination of 5%. In dereplication, we implemented a primary clustering threshold (-pa) of 0.9 and secondary alignment threshold (`-sa`) of 0.95, requiring minimum overlap between genomes (`-nc`) of 0.3, using multi-round primary clustering (`--multiround_primary_clustering`) and greedy secondary clustering with fastANI v1.33^96^ (`--greedy_secondary_clustering`, `--S_algorithm fastANI`) to reduce the computational complexity of dereplicating a large genome set. For dereplication, cluster representatives were chosen using scoring criteria that included a completion weight (`-comW`) of 1, contamination weight (`-conW`) of 5, N50 weight (`-N50W`) of 0.5, size weight (`-sizeW`) of 0, and centrality weight (`-centW`) of 0. Genome filters and scoring were consistent with standards used in the UHGG^49^. The final genome set was taxonomically classified and placed in a tree with GTDB-tk v2.3.0^97^ using the GTDB r214 catalogue and default parameters. Phylogenetic trees were visualized with iTOL v6^98^. The dereplicated prokaryotic genome set was compared against the UHGG v2.0.1 species representatives using dRep v3.4.3 with the same parameters as previously stated above.

Protein-coding genes were predicted from each medium-quality and high-quality prokaryotic genome, prior to genome dereplication, using prodigal v2.6.3 with parameters `-c -p meta` to exclude partial genes. Putative proteins were clustered successively using mmseqs v14.7e284^99^ linclust command with alignment coverage (-c) of 0.8 in target coverage mode (`--cov-mode 1`) and greedy secondary clustering (`--cluster-mode 2`) at 100% and 95% amino acid identity (`--min-seq-id`). The 95% identity protein set was compared against the UHGP95 v2.0.1 proteins using mmseqs v14.7e284, and proteins sharing 95% amino acid identity over 80% of the UHGG protein were considered to match the UHGP set.

Viral genomes were annotated with VIBRANT v1.2.1^84^ and genome quality was determined with checkV v1.0.1^100^. Viral genomes were clustered into a dereplicated set by building a database of medium- and high-quality genomes with BLAST 2.14.0^101^ and clustering genomes at a minimum 95% ANI and 85% alignment fraction using checkV supporting scripts with default parameters. The final viral genome set was compared against the MGV v1.0^102^ vOTU representative phage genomes using the same BLAST clustering approach and parameters.

Modelled accumulation of novel prokaryotic genomes, proteins, and viral genomes with additional participant sampling was determined by randomly subsetting the full participant set or site-specific participant sets to a range of individuals (1-1500) in a hundred iterations and counting the number of prokaryotic genome clusters, protein clusters, or viral genome clusters represented by the participant subset.

### *Treponema succinifaciens* core genome determination and phylogeography analysis

We evaluated the complete set of *T. succinifaciens* metagenome-assembled genomes in our genome catalogue prior to dereplication. To identify *T. succinfaciens* genomes, we selected all genomes with completeness ≥50% and contamination ≤5% that fell into a secondary cluster with genomes classified as *Treponema_D succinifaciens* by GTDB-tk in our dereplicated genome catalogue (*n* = 249). Coding sequences were predicted with bakta v1.8.2^103^. Core genes, defined here as genes present in at least 80% of genomes, were identified with roary v3.12.0^104^.

Public *T. succinifaciens* genomes were downloaded from the UHGG^49^, Carter *et al.*^50^, and NCBI. To build AWI-Gen specific or global phylogenetic trees, core genes were identified and incorporated into a core gene multiple sequence alignment using roary v3.12.0^104^ and MAFFT v7.407^105^. The core gene multiple sequence alignment was used as input to FastTree v2.1.11^106^, and the resulting phylogenetic trees were visualized in iTOL v6^98^.

### Association between microbiome features and HIV status

Participants from Agincourt, South Africa, Soweto, South Africa, and Nairobi, Kenya were included in this analysis. Participants from DIMAMO, South Africa were excluded due to the low number of PLWH (*n*=6). Participants from Nanoro, Burkina Faso and Navrongo, Ghana were excluded as HIV status was not measured in these populations due to low national prevalence of HIV. A total of 850 participants was included in this analysis, capturing 129 people living with HIV (PLWH) and 721 seronegative individuals (see **Table 1**). The rest of the samples from those sites were either HIV positive, but reported not to take antiretroviral therapy (*n*=28, *n*=22 in Agincourt, *n*=3 in Soweto, *n*=3 in Nairobi), or information about ART or HIV status was missing. Prokaryotic alpha and beta diversity were calculated as described above.

Differential abundance analysis was performed using a linear mixed effect model implemented in the lmerTest R package v3.1-3^107^, including site, exposure to antibiotics, and self-reported recency of diarrhoea as random effects, as those factors had shown to be related to microbiome composition in the previous analyses. Overall effect size was estimated through the lmerTest package as well and generalized fold change within each site was calculated as described above.

For the machine learning analysis, we trained statistical models using the SIAMCAT R package v2.5.0^86^ for both all data combined and for each site separately. In short, relative abundances were normalized using the `log.std` method in SIAMCAT. Samples were split for five-times repeated five-fold cross-validation (20% of samples were retained for testing and not included in model training) and for each split, an L1-regularized logistic regression model (LASSO) was trained on the training folds, using standard parameters. Model evaluation was performed within the cross-validation (for example, within a site) by applying each model to the respective left-out test fold. The predictions for each sample were averaged across repeats and the area under the receiver operating characteristics (AU-ROC) was calculated with the pROC package v1.18.2^108^. For cross-site evaluation, the external data was normalized with the recorded normalization parameters (frozen normalization), all models from the cross-validation were applied to the normalized data, and predictions were averaged again for AU-ROC analysis. For the leave-one-site-out (LOSO) analysis, models were trained as described on data from two sites combined (for example, Agincourt and Soweto) and were then applied on the data from the left out site (Nairobi).

To test the fraction of positive prediction in other sites, we calibrated the model prediction to an internal 5% false positive rate, that is, recorded at which prediction threshold 5% of HIV-samples were incorrectly classified as PLWH. Then, the model trained on all data combined was applied to the data from Nanoro, Navrongo, and DIMAMO to quantify the number of samples that resulted in a prediction above the threshold value.

### Statistical analysis

Statistical analyses were performed using R v4.1.2 using the statistical test specified in the respective methods section. Correction for multiple hypothesis testing was performed with the Benjamini-Hochberg procedure^109^ as implemented in the `p.adjust` function in base R. Plots were generated in R using the following packages: ggplot2 v3.4.2^110^, cowplot v1.1.1^111^, tidyverse v2.0.0^112^.

## Data Availability

All shotgun sequence data and the dereplicated prokaryotic and viral genome sets will be deposited in the European Nucleotide Archive upon publication. Participant phenotype data will be deposited in the European Genome-Phenome Archive upon publication. Participant phenotype data are under restricted access due to ethics requirements of the AWI-Gen 2 study. Access can be obtained by request from the Human Heredity and Health in Africa Data Access Committee at https://catalog.h3africa.org/. Source data, including classification tables, genome summary statistics, taxon prevalence, and differential feature tables, are available at https://purl.stanford.edu/kq420ys8307.

Reference data used in this study are available from the Unified Human Gastrointestinal Genome collection in the European Nucleotide Archive under project accession PRJEB33885, the Metagenomic Gut Virus catalogue at https://portal.nersc.gov/MGV, and the Genome Taxonomy Database at https://data.gtdb.ecogenomic.org/releases/.

## Code Availability

Nextflow workflows for metagenomic preprocessing, assembly, binning, and postprocessing, as well as the source code for analysis and figure generation are available at https://github.com/bhattlab/AWIGen2Microbiome.

## Acknowledgements

We thank all of the study participants for participating in this research project, along with each community advisory group for their recommendations on study design. We thank the field workers and staff at each study centre for participant enrollment and sample collection. We thank Navrongo centre PI Dr. Abraham Oduro and NHRC Director Dr. Patrick Ansah, as well as Caeser Santuah and Rita Afiya for supporting sample collection. We thank the Soweto study and data collection team: Vukosi Baloyi, Sphe Sekwati, Thonniah Ngobeni, Onke Godongwana, and Michael Ritsuri. We thank Cassandra Soo for her role as biobank manager through 2021 and for assistance with standard operating procedure development and sample shipping. We thank the Stanford Research Computing Center for computational infrastructure. We also thank Erin Brooks, Gabriella Reynolds, and Carrie Chen for assistance in sample processing, Stephen Nayfach for advice regarding phylogeography calculations, and Michele Barry and Steve Luby for guidance regarding the project.

This work was supported by the National Human Genome Research Institute (NHGRI), the Eunice Kennedy Shriver National Institute of Child Health & Human Development (NICHD) and Office of the Director (OD) of the National Institutes of Health (NIH) of the USA under award number U54HG006938 for the AWI-Gen study, and its supplements, as part of the H3Africa Consortium. Additional funding was granted by the Department of Science and Technology (now Department of Science and Innovation), South Africa (award number: DST/CON 0056/2014). This work was also supported, in part, by the Stanford Center for Innovation in Global Health. This work used supercomputing resources provided by the Stanford Genetics Bioinformatics Service Center, supported by NIH S10 Instrumentation Grant S10OD023452. D.G.M. was supported by the Stanford Gerald J. Lieberman Fellowship and the NIH Fogarty Global Health Equity Scholars Program (NIH FIC D43TW010540). J.W. acknowledges support from the School of Medicine Dean’s Postdoctoral Fellowship. V.O. was partially supported by the NIH Fogarty Global Health Equity Scholars Program (NIH FIC D43TW010540). M.R. is the South African Research Chair in Genomics and Bioinformatics of African Populations hosted by the University of the Witwatersrand (SARChI), funded by the South African Department of Science and Innovation, and administered by the National Research Foundation. The Bhatt lab is supported by NIH R01AI148623 and R01AI143757, a Stand Up 2 Cancer Grant. A.S.B is supported by the Allen Distinguished Investigator Award. S.H. was partially supported by the SA National Research Foundation (CPRR160421162721).

This paper describes the views of the authors and does not necessarily represent the official views of the National Institutes of Health (USA), the South African Department of Science and Technology or the National Research Foundation (South Africa) who funded this research.

## Author Contributions

A.S.B., S.H., M.R. conceived of the study, secured funding, and obtained necessary ethics approvals. S.H., O.H.O., M.R., F.T., G.A., P.R.B., S.S.R.C., F.X.G.O., I.K., G.R.M., L.M., S.F.M., E.A.N., S.N., H.S., S.T., and F.W. organized study logistics, community engagement, participant enrollment, and data collection. N.S., O.H.O., and D.G.M. coordinated sample preparation, shipping, and sequencing. D.G.M., J.W., L.A.I.O., T.M., and C.W.B. performed data analysis. D.G.M., O.H.O., J.W., L.A.I.O., A.S.B., and S.H. wrote and edited the manuscript. A.S.B. and S.H. supervised the work and oversaw the collaboration. All authors have read and approved the manuscript.

## Ethics and Inclusion Statement

All authors of this study fulfilled criteria for authorship inclusion, and researchers from each study centre are represented as authors. Researchers from all institutions were involved throughout the study process. Study centre staff facilitated community engagement sessions, which identified specific community concerns and determined that this study is locally relevant. Roles and responsibilities were agreed upon amongst collaborators prior to conducting the research. Authors of this study have led formal capacity-building genomics workshops for local scientists during the course of the study (see Figure 1), along with additional informal training. This study has been approved by local ethics review committees (see Methods). Research pertinent to the study centres and led by local researchers has been taken into account in the citations.

## Additional Information

Supplementary Information is available for this paper. Correspondence and requests for materials should be addressed to Ami S. Bhatt or Scott Hazelhurst.

