## Supplementary Methods for "Expanding the human gut microbiome atlas of Africa"

### Cohort description and participant selection

The AWI-Gen project is a study of genomic and environmental risk factors for cardiometabolic disease in population cross-sectional cohorts from six study sites across Africa. Five of the six study sites are managed by health and demographic surveillance systems (HDSSs) that monitor regional population health. Each HDSS maintains detailed residency, migration, and other census information for all individuals in their catchment areas. Leveraging this census and residency information, the AWI-Gen was able to randomly enrol participants from across each study site after filtering the potential participant pool based on inclusion and exclusion criteria.

Phase 1 of AWI-Gen was conducted between 2012 and 2017 and recruited ~10,700 individuals 40–60 and ~1,200 individuals 60-74[1]. Phase 2 of AWI-Gen followed up this cohort in 2018-2022. All AWI-Gen 1 participants were eligible for AWI-Gen 2, but due to death, out-migrations, unavailability, and refusals to participate, only 7,225 individuals (4,124 women) participated in AWI-Gen 2. The DIMAMO site recruited a small number of new participants, but most AWI-Gen 2 participants were in also AWI-Gen 1.

*Recruitment approach*

Our earlier work[2,3] was a pilot study of ~200 AWI-Gen 1 participants, intended to determine feasibility of incorporating a microbiome component into AWI-Gen and perform preliminary analyses of microbiome composition and obesity in two South African populations. In this paper, we report on the microbiome component of AWI-Gen 2. Due to budgetary constraints, we had to sub-sample participants from the overall AWI-Gen 2 study. Therefore, we semi-randomly chose 1,825 participants from the AWI-Gen 2 female participants. We aimed at recruiting women, primarily because menopause and hormonal transition is another arm of the overall AWI-Gen 2 study and we want to explore the link between the microbiome and menopause in subsequent research. We enriched for recruitment of younger women who were most likely to still be pre-menopausal in AWI-Gen 2 based on data from AWI-Gen 1. An error at one of the sites led to a small number of men being recruited into the study. Since Kenya and South Africa have a high proportion of people living with HIV, and we wanted to reduce the confounding factors of HIV and antiretroviral treatment, we decided to focus our study on HIV– individuals. However, at recruitment we only had partial knowledge of HIV status. Rapid tests were undertaken at point of collection, but for ethical reasons we decided that any participant who volunteered to participate in the microbiome study would be enrolled, since otherwise their HIV status would be obvious to many field workers who should not be privy to this information. We also recognized that a small sub-cohort of HIV+ participants would be a valuable addition, considering the small number of microbiome studies in African populations with individuals who are HIV+.

*Recruitment methodology*

We pursued an ambitious plan for subsampling AWI-Gen 2 participants for participation in the microbiome sub-study. Our sampling approach is described below. Each site received a ranked list of participants for inclusion into the microbiome study, with the request that they recruit accordingly but diverge as needed. Due to complicated sampling logistics, the following lists were not strictly adhered to.

Although this is a cross-sectional study, we particularly wanted to ensure that the study population would be sufficient for future studies into menopause and hypertension, as these conditions are of particular interest to the overall AWI-Gen 2 study. We enriched our sampling for women in different categories rather than attempting to specifically build case-control cohorts. The logistics of a large-scale cross-sectional study imposed restrictions, especially as we needed to reduce the burden on field-workers and participants by reducing the number of visits. Therefore, we created a ranked list of all women at each site as follows:

- A1: a random selection of women who were pre- or peri-menopausal and not hypertensive in AWI-Gen 1;
- A2: a random selection of women who were pre- or peri-menopausal and hypertensive in AWI-Gen 1;
- B1: a random selection of women who were post-menopausal and not hypertensive in AWI-Gen 1;
- B2: a random selection of women who were post-menopausal and hypertensive in AWI-Gen 1;
- C: all other women, randomly ordered.

Participants randomly selected into the *A* and *B* lists were prioritized for recruitment. As we anticipated loss-to-follow up, we randomized more participants (by 10%) into these lists than our target total number of participants per site. Additionally, since 5 years had elapsed since AWI-Gen 1 and many of the women who had not been post-menopausal would have become post-menopausal in AWI-Gen 2, we also prioritized enrolment of participants from the pre-menopausal *A* lists (by an *A*:*B* ratio of 3:2). We asked each site to try to recruit the women in the microbiome sub-study in order of the ranked list. However, as explained above, recruitment into the microbiome sub-study was embedded into the broader AWI-Gen 2 study, and the complexity of large scale recruitment meant that it was not always possible to adhere to the priority list. Sites were able to enrol participants from list *C* where it made logistical sense, or due to unavailability of higher-ranked participants. Finally in the Agincourt site, we tried to recruit the small number of women who were second degree or closer relatives (based on the genetic data from AWI-Gen 1). Note that at the point of recruitment into AWI-Gen 1, there was only one participant per household.

*Enrolment statistics*

Loss to follow-up from AWI-Gen 1 to was roughly 35%. There were a significant number of refusals of individuals from AWI-Gen 1 to participate in the Phase 2 study, and this varied from site to site, but to our knowledge no person refused on account of the microbiome study. It was possible for participants to selectively refuse to participate in the microbiome study and continue with the rest of the AWI-Gen study.

The table below summarizes the characteristics of our participants and all women in AWI-Gen 2. Except for modest boosting of the number of pre-menopausal women, participants in the microbiome component match the overall AWI-Gen 2 participants

| | **Microbiome participants** | **All women** |
|---|---|---|
| **Mean BMI** | 27.4 | 27.5 |
| **Mean age** | 55.2 | 57.1 |
| **Mean systolic blood pressure** | 128.2 | 127.5 |
| **% pre-menopausal (AWI-Gen 2 data)** | 13.7 | 9.4 |

The table below shows the number of participants in the microbiome study compared to the total number of women participating in AWI-Gen 2. We had a larger budget for recruitment at Agincourt and Nanoro, hence the larger numbers there.

| **Site** | **Agincourt** | **DIMAMO** | **Nairobi** | **Nanoro** | **Navrongo** | **Soweto** |
|---|---|---|---|---|---|---|
| Women in AWI-Gen 2 | 660 | 768 | 694 | 784 | 536 | 705 |
| In microbiome sub-study | 533 | 203 | 237 | 382 | 235 | 226 |

### Site descriptions

This section supplements the overview of the sites shown in **Figure 1**. Site characteristics are described below, and a selection of key statistics are in **Table 1**. The six AWI-Gen sites are all active research sites involved in ongoing health and demographic surveillance. The AWI-Gen project was one of the major projects over the 2012-2023 period.

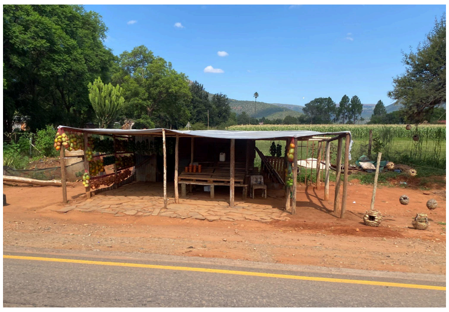

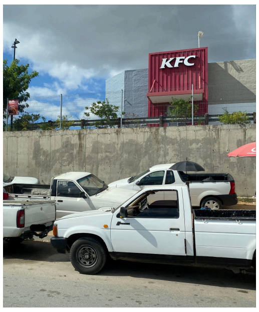

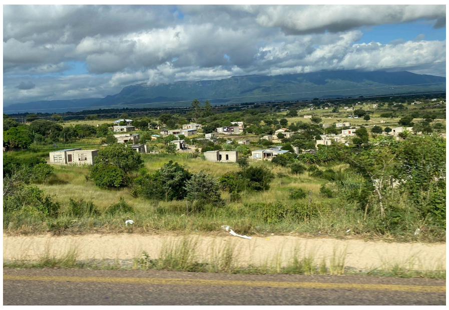

**Agincourt, South Africa:** The Agincourt HDSS (named after the small village of Agincourt, Mpumalanga where the HDSS unit's offices and labs are) was established in 1992 in the Bushbuckridge sub-district of Mpumalanga Province in South Africa[4,5]. The district has a complex political and social history that has shaped population growth and migrancy over time: during apartheid (1948-1990) the policy of forced removal brought thousands of people into the district at different times from different places, some of whom had no previous link to the area. The Mozambican civil war in the 1980s led to the arrival of significant numbers of refugees who integrated into the community[4,6]. The district therefore has a complex cultural and linguistic mix. When the HDSS was established in 1992, Bushbuckridge was a very rural site. It has since undergone significant epidemiological transition. Although the population density remains very low compared to the urban sites, shopping centres and fast food outlets now exist in the district. The government has put considerable effort into reticulation of electricity and water, but the use of fuels other than electricity for cooking is common, and piped water distribution remains a challenge. Over half of the houses have toilets outside of the home. The surveillance site now consists of 27 villages (political entities)/31 villages (observational units).

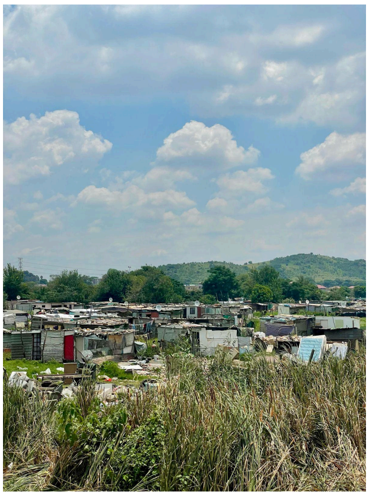

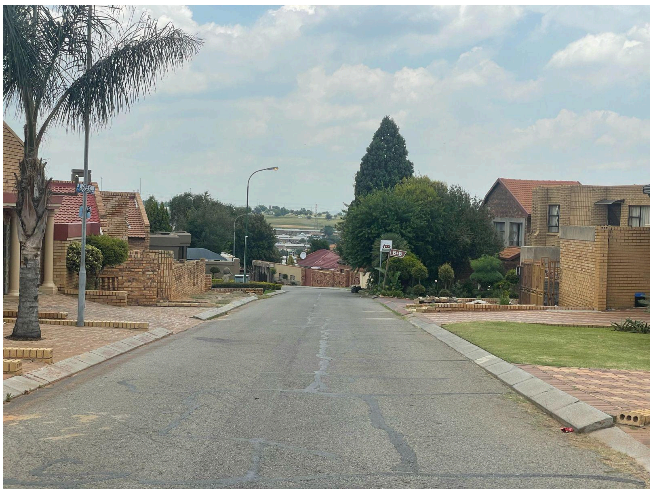

**Soweto, South Africa**: Soweto, an area of Johannesburg, was established between 1930 and the 1950s for black residents of Johannesburg, and many of the initial residents were forcibly moved from other urban areas in Johannesburg. Although Soweto has always formally been part of the Johannesburg geographical area, it was only in 1994 that the city had a unitary government. The complex administrative history has led to widely varying estimates of population. The 2011 census estimated Soweto's population at 1.2 million (of the 4.4 million in Johannesburg, with an estimate that 13.1 million people lived within a radius of 100 km of Johannesburg). Since 2011, a population growth of 20% is likely[7]. Soweto is arguably the most diverse of our six sites with significant socio-economic differentiation[8]. There are areas of informal settlement where people live in shacks but also very middle-class areas. Poverty levels remain high. In 2016, 94.1% of Johannesburg residents had access to safe drinking water, and 88.6% had flush toilets[9]. More granular results for Soweto are not available. The proportion of individuals with access to safe drinking water and flush toilets is likely to be lower in Soweto than the average in Johannesburg, but still high compared to other sites.

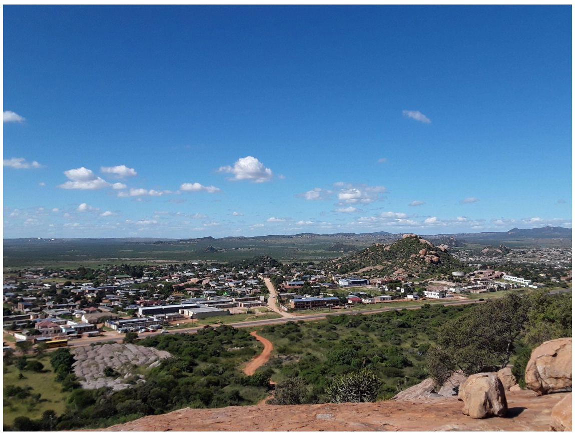

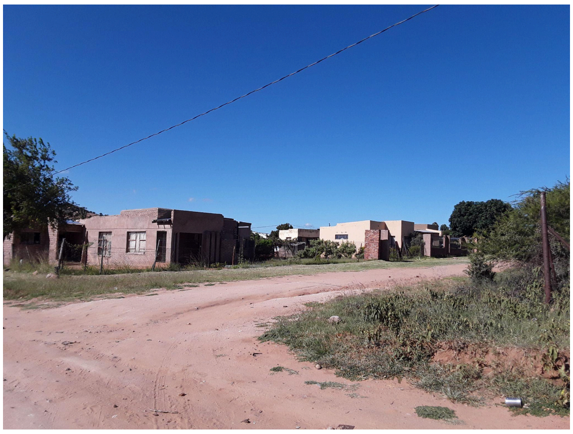

**DIMAMO, South Africa:** The Dikgale HDSS was established in 1995 with the aim of collecting data relating to migration, socioeconomic status, cardiovascular risk factors and household global positioning system (GPS) coordinates among other variables[10]. The surveillance area was later expanded to include villages that form part of the Dikgale and Mamabolo tribal areas, hence the name change to DIMAMO Population Health Research Centre (PHRC). The HDSS is located in the Capricorn District of Limpopo Province, South Africa and captures 57 rural and semi-urban villages. Black people make up a majority of the population, and Northern Sotho is the dominant language. The greater part of the population is of low economic status and educational levels, with high levels of unemployment[11]. Some members of the community are involved in subsistence farming, with agriculture constituting the primary subsistence type. Although the DIMAMO catchment area is mostly rural, the villages have access to piped water and electricity; however, households still utilize other sources of energy such as firewood and gas. The surveillance region is serviced by 11 primary health care facilities and one tertiary hospital.

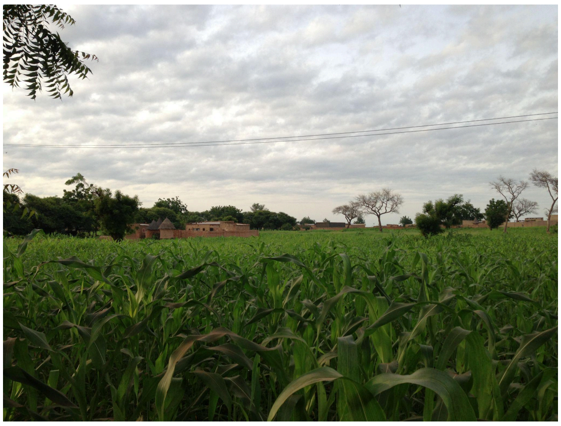 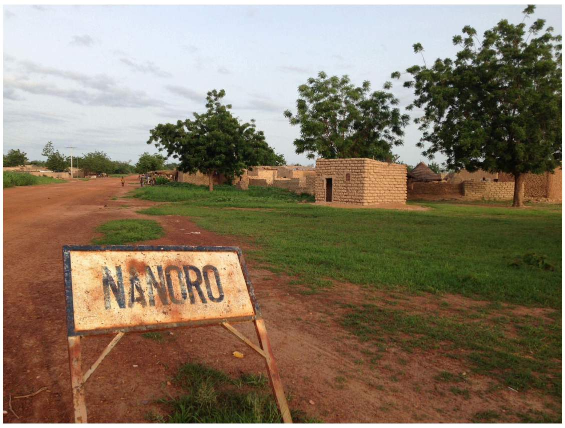

**Nanoro, Burkina Faso:** The Nanoro HDSS was established in 2009 by the Clinical Research Unit of Nanoro (CRUN)[12]. Nanoro is in central Burkina Faso, approximately 85 km from Ouagadougou. The demographic surveillance area lies within the Health District of Nanoro and covers 594.3 km$^2$, about ~36% of the total Nanoro Health District area. The HDSS serves as a platform for clinical research and training on diseases of local public health importance, and the current research portfolio includes malaria, other infectious diseases, chronic diseases and genomics, and impact of climate change on health. Most of the population practices agriculture, primarily subsistence farming and horticulture.

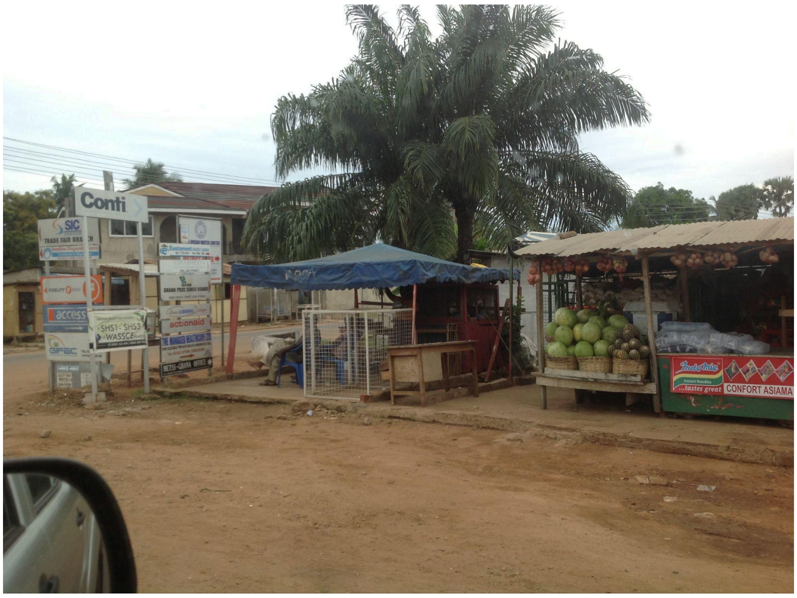

**Navrongo, Ghana:** The Navrongo Health Research Centre (NHRC) in northern Ghana started in 1988 as a field site for vitamin A supplementation trial through collaboration between the London School of Hygiene and Tropical Medicine (LSHTM) and the Kwame Nkrumah University of Science of Technology (KNUST) with the support of the Ministry of Health. In 1992, the Navrongo Health and Demographic Surveillance System (NHDSS) was established to collect data on births, deaths, migrations, socioeconomic status among others[13]. The Navrongo Health Research Centre conducts studies including clinical trials, social and demographic research, communicable and non-communicable diseases research. The study area lies between latitude 10.30’ and 11.10’ north and longitude 1.1’ west, and covers a total land area of 1675 km$^2$, bordering northwards along the

Ghana-Burkina Faso border. The area is semi-arid and primarily covered by Guinea savannah vegetation consisting of grassland integrated with short trees. The NHRC collects information from over 157,000 people from two districts (Kassena-Nankana Municipality, and Kassena-Nankana West) in northern Ghana using the HDSS[13]. Over 70% of people within the site are subsistence farmers, and the remainder of the population is in public sector employment or unemployed. About 69.1% of the population within the NHDSS catchment area have access to electricity. Access to piped water is about 15.9% while access to borehole water is 81.6%. The rest drink from closed and open wells. About 23.6% of the people in the site have access to toilet facilities. People within the HDSS site use various methods of cooking with the use of firewood being the comments method (70.4%). This is followed by charcoal (19.2%) and gas (7.4%) with the rest using other methods.

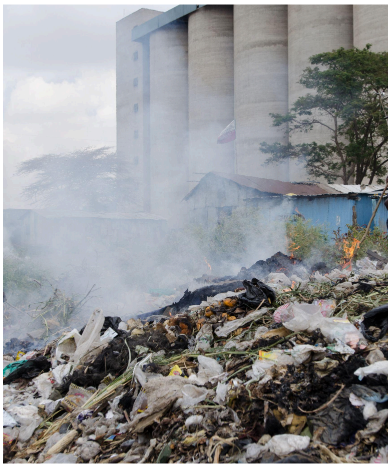

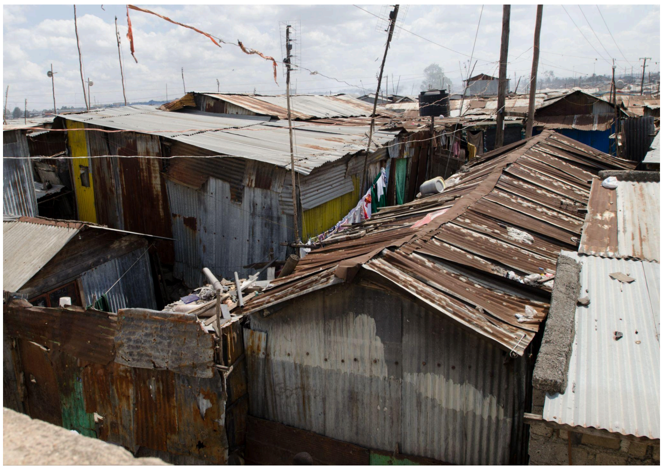

**Nairobi, Kenya:** The Nairobi Urban Health and Demographic Surveillance System (NUHDSS) was established in 2002 to provide research infrastructure to study the health consequences of urban residence, and to evaluate the impact of intervention studies focused on the urban poor[14,15]. The NUHDSS captures the two urban informal settlements of Korogocho and Viwandani, both located in Nairobi, Kenya. The NUHDSS follows a population of about 65,000 individuals and collects key demographic and health information and data related to the living conditions in the informal settlements, such as employment opportunities and household amenities. Viwandani borders a nearby industrial area; most residents of Viwandani are younger, mobile, and actively working or seeking jobs. The population of Korogocho is more stable, and many residents have lived in the area for several years[14]. Korogocho has the worst health and socio-economic outcomes of all the informal settlements in Nairobi, with limited access to education, employment, water and sanitation[14,16].

### Participant covariate processing

Extensive participant data was collected as part of the AWI-Gen study, including demographic, ethnolinguistic, family composition, pregnancy, cognition, frailty, household amenity, substance use, general health, diet, infection history, cardiometabolic disease, and physical activity information. Participants also gave blood, urine, and stool samples, and underwent ultrasound, blood pressure, blood, and urine testing for various metrics. Not all data were available for every participant, and some participants gave stool samples for microbiome analysis but did not complete other testing or questionnaires. At the time of analysis for the microbiome study, not all participant data had gone through quality control. In total, 59 variables were available to use as covariates in the microbiome study.

Prior to using covariate data in microbiome analysis, we first collapsed the covariate dataset to only those variables that we expected to be most meaningful to avoid unnecessary multiple-hypothesis testing and measuring associations between dependent variables. First, we removed variables that had overwhelmingly missing data, excluding those that had entries for 100 or fewer participants (e.g. several ultrasound measurements). Second, we filtered variables with not enough unique values (such as sex, which had only one group). Lastly, we excluded variables with an entropy (calculated with the `infotheo` package v1.2.0.1[17] in R) of less than 0.2 to avoid variables that were too uniform in the participant set to power comparisons (e.g. breast cancer or cervical cancer status with only 10 and 12 cases, respectively).

To calculate correlation between covariates and associations between covariates and microbiome composition, we transformed non-numerical covariates into numerical values based on ordered factor levels. For example, values for the *Menopause* covariate were changed from `Pre-menopausal` to 1, from `Peri-menopausal` to 2, and from `Post-menopausal` to 3.

Most covariates were binary (for example, *Probiotics* could contain either the value `Yes` or `No`) and were converted to 1 (for `Yes`) and 2 (for `No`) in this process. The full list of binary variables is: *Arthritis*, *Diabetes status*, *Diabetes treatment*, *Hypertension status*, *Hypertension treatment*, *Pesticides*, *Vigorous work*, *Weekend work*, *HIV medication*, *HIV status*, *Cattle*, *Other livestock*, *Potable water*, *Poultry*, *Refrigerator*, *Toile*t, *Deworming treatment*, *Probiotics*, *Chew tobacco* and *Smokeless tobacco*. The variables describing time periods (*Deworming period*, *Probiotics period*, *Antibiotics*, and *Diarrhoea last*) were ordered according to recency with this order:

- `WithinLastWeek` < `WithinLastMonth` < `WithinLastSixMonths` < `WithinLastYear` < `WithinLastTwoYears` < `WithinLastThreeYears` < `Longer` < `Never`.

All other variables were converted as follows: *Menopause* see above, *Employment* was ordered as

- `Self-Employed < FormalFull-time < FormalPart-time < Informal < Unemployed`,

and Site was ordered as

- `Nanoro` < `Navrongo` < `Dimamo` < `Agincourt` < `Soweto` < `Nairobi`.
