## Supplementary Figures for "Expanding the human gut microbiome atlas of Africa"

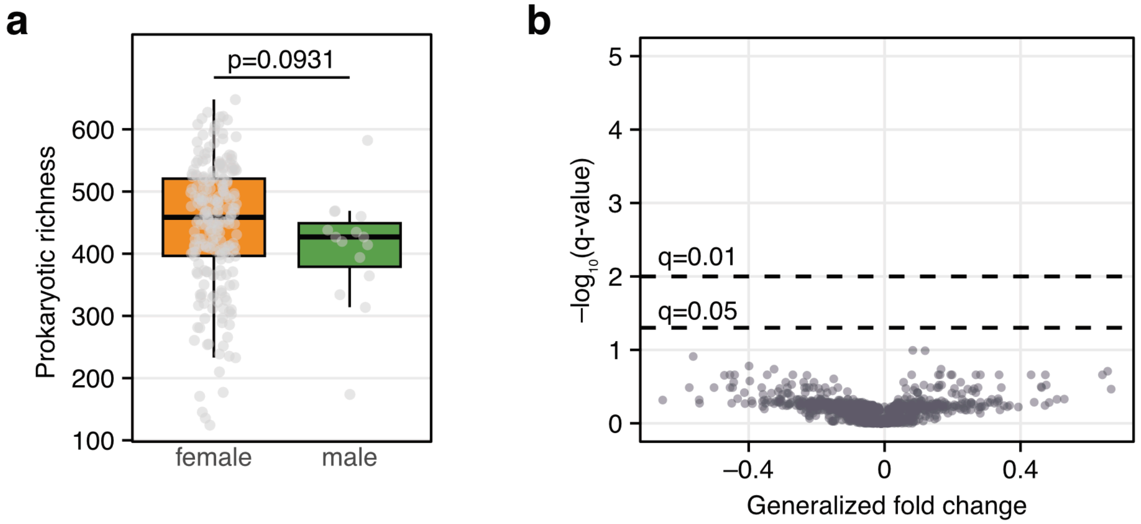


**Supplementary Figure 1. Microbiome composition of male and female participants in Navrongo, Ghana. a)** Prokaryotic richness (number of prokaryotic species present at ≥1e-04% abundance after rarefaction, see Methods) in n = 16 males and n = 218 females in Navrongo, Ghana (Wilcoxon test, p = 0.0931). Points indicate individual samples. (In total, 17 samples from male participants were sequenced, but the missing male sample corresponds to a single male participant from Agincourt; this sample was excluded for this analysis to account for differences between sites.) **b)** Generalized fold change between male and female participants for all species with a prevalence higher than 5% in Navrongo is plotted against the negative log10-transformed q-value (Benjamini-Hochberg corrected p-value). Positive values correspond to higher relative abundance in males, whereas negative fold change values indicate higher relative abundance in female participants. No species meet the threshold of significance after correction for multiple testing. For all boxplots, boxes denote the interquartile range (IQR) with the median as a thick black line and the whiskers extending up to the most extreme points within 1.5-fold IQR.

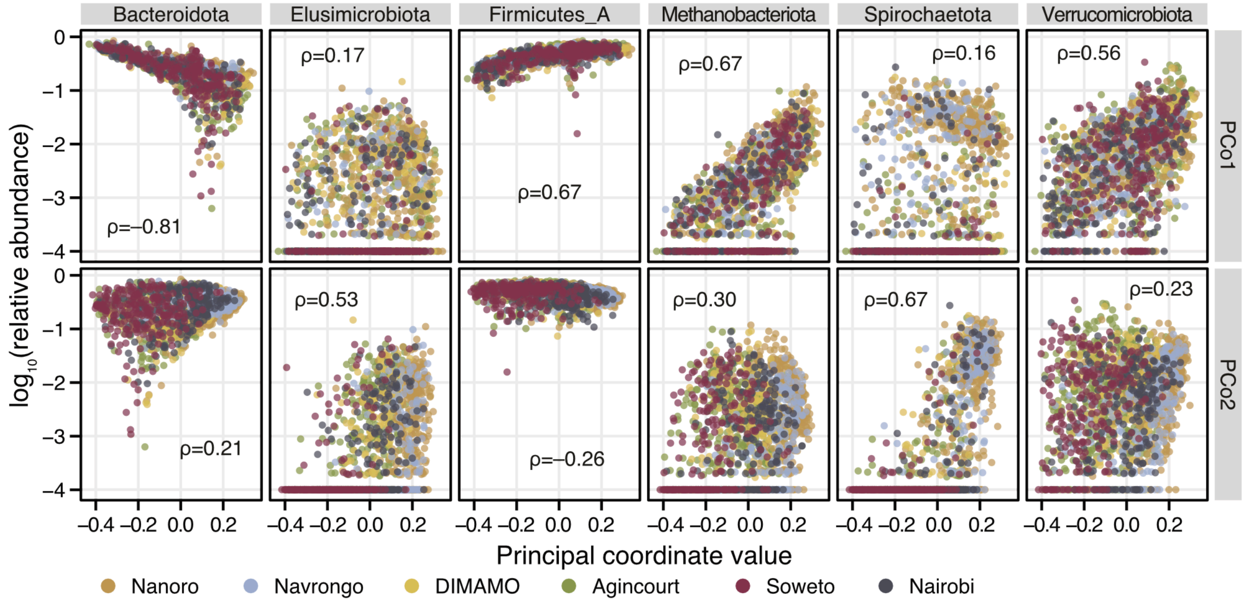


**Supplementary Figure 2. Correlation between principal coordinate values and prokaryotic phyla.** Spearman correlation coefficient (Spearman's rho) between principal coordinate values and the relative abundance of selected prokaryotic phyla. Phyla with an absolute correlation coefficient higher than 0.4 for either of the first two principal coordinates are shown (see **Figure 2** in the main text). Points represent individual samples and are coloured by site.

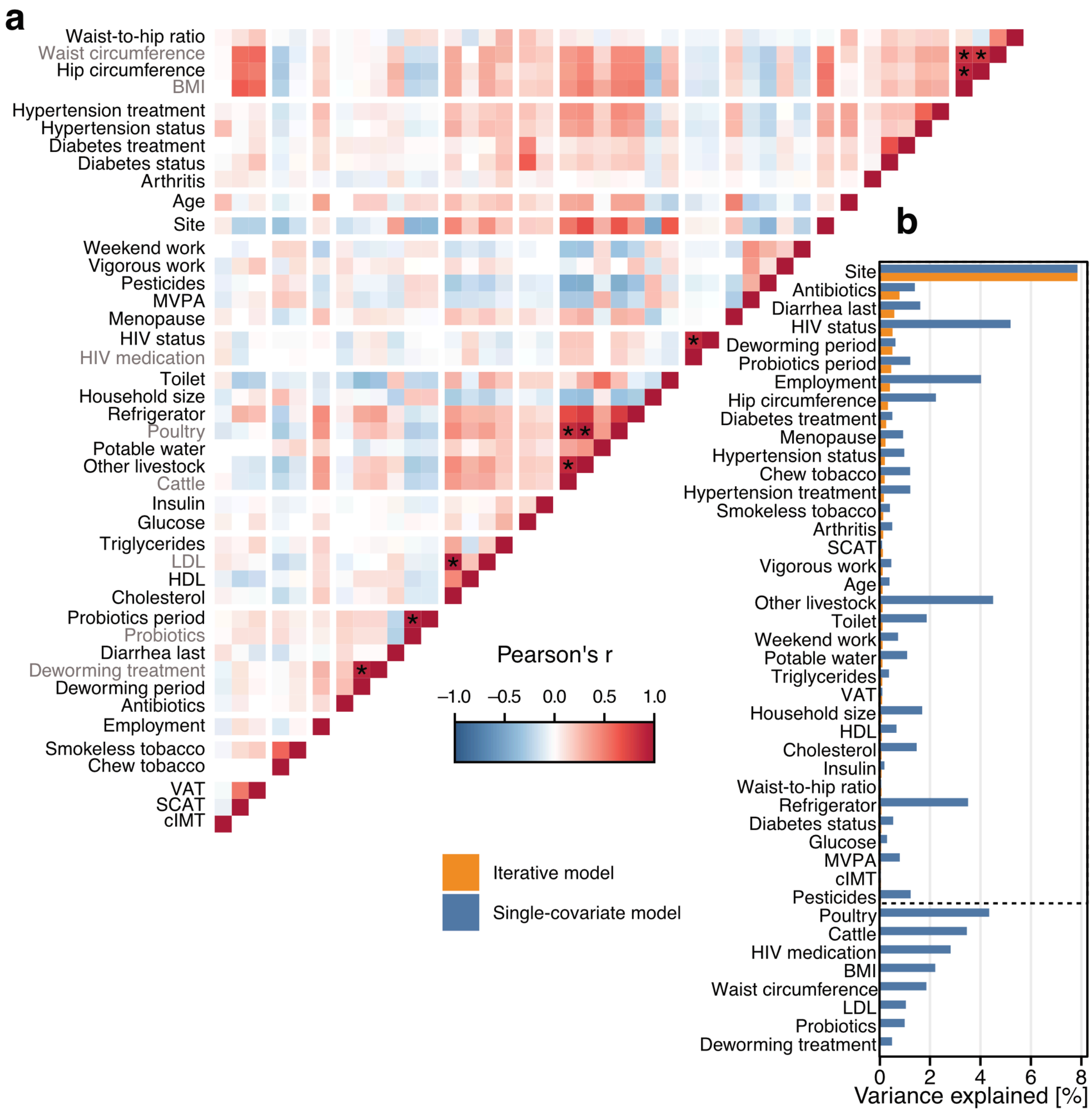


**Supplementary Figure 3. Metadata correlation and distance-based redundancy analysis. a)** Pearson correlation coefficient (Pearson's r) between available participant covariates, calculated on all participants included in the site comparison (*n*=1796). Non-numerical covariates were transformed into numerical values based on ordered factor levels (see Supplementary Methods). Asterisks indicate highly correlated covariates (Pearson's r ≥ 0.8). In those cases, the covariate that explained the higher amount of variance in the prokaryotic composition (see panel b) was selected (redundant variables are indicated by grey labels). **b)** The amount of variance in the prokaryotic composition that is explained by covariates in distance-based redundancy analysis. Blue bars indicate single-covariate models (each covariate associated with prokaryotic composition individually), whereas orange bars show the amount of variance explained in the iterative model in which the variable explaining the most additional variation is added iteratively to a multi-covariate model (see Methods). Covariates below the dashed line were removed before the iterative modelling since they were highly correlated with other covariates. BMI: body mass index, MVPA: moderate to vigorous physical activity, LDL: low-density lipoproteins, HDL: high-density lipoproteins, VAT: visceral adipose tissue, SCAT: subcutaneous adipose tissue, cIMT: carotid intima-media thickness.

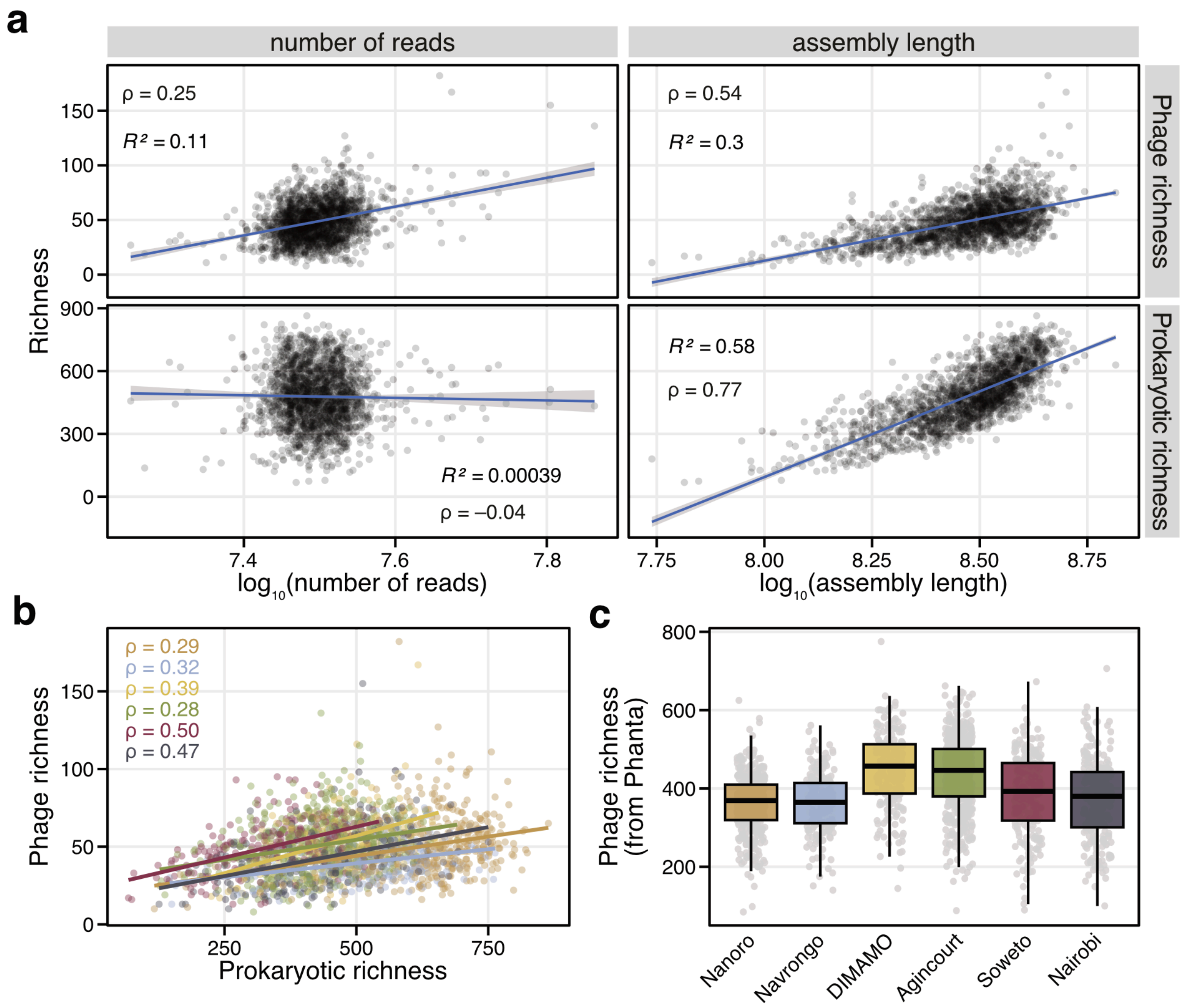


**Supplementary Figure 4. Phage and prokaryotic alpha diversity. a)** Spearman correlation coefficient (Spearman's rho) and $R^2$ for a linear model between phage richness (number of assembled phages) or prokaryotic richness (number of prokaryotic species present at ≥1e-04% relative abundance after rarefaction) and total read count and total assembly length (length of the total assembly in base pairs). Points represent individual samples. Blue line indicates a linear association model with 95% confidence intervals shown as shaded areas. **b)** Spearman correlation between prokaryotic richness and phage richess. Points represent individual samples and are coloured by site (see panel c) for a colour-code). **c)** Phage richness per sample, based on Phanta profiles (number of phage species clusters present ≥1e-04% relative abundance). For all boxplots, boxes denote the interquartile range (IQR) with the median as a thick black line and the whiskers extending up to the most extreme points within 1.5-fold IQR.

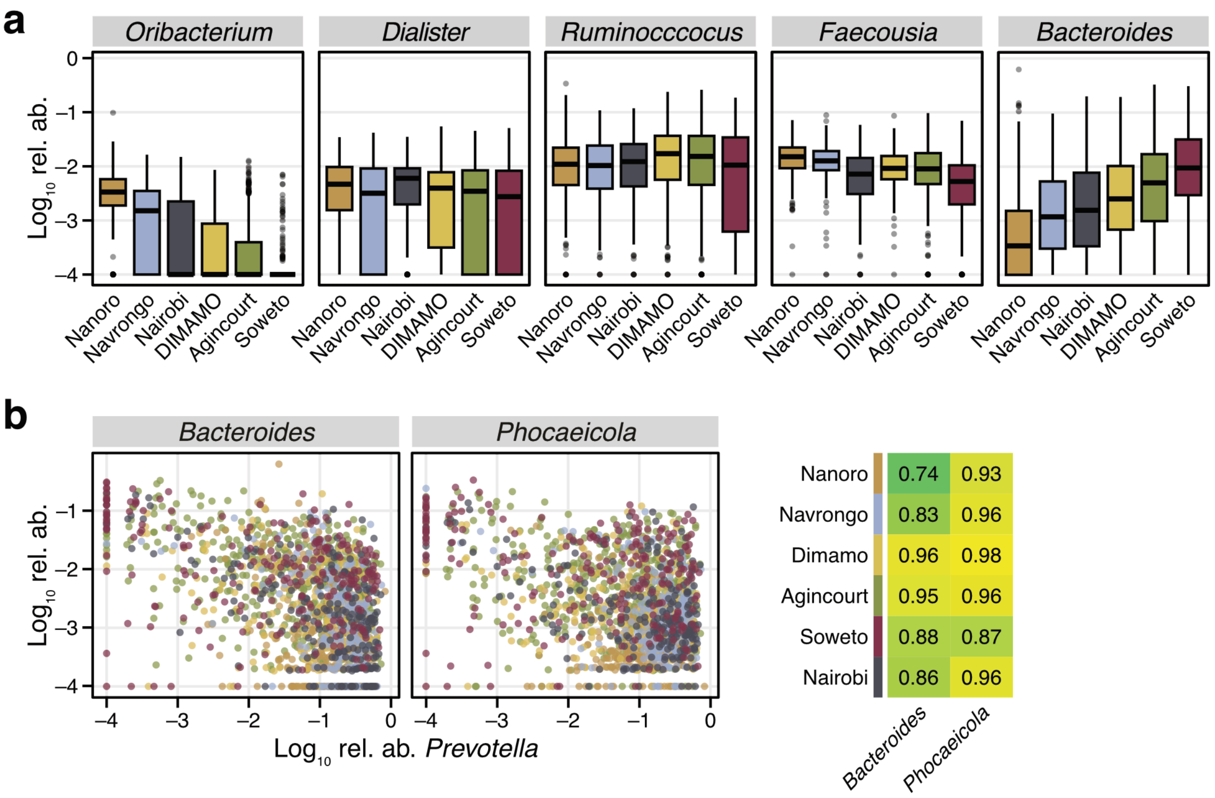


**Supplementary Figure 5. Relative abundance of prokaryotic genera. a)** The log10-transformed relative abundance of select genera are shown across the six different study sites (sites ordered by the clustering in Fig. 2 of the main text). For all boxplots, boxes denote the interquartile range (IQR) with the median as a thick black line and the whiskers extending up to the most extreme points within 1.5-fold IQR. **b**) The log10-transformed relative abundance of the genus *Prevotella* plotted against the relative abundance for the genera *Bacteroides* and *Phocaeicola*. Points represent individual samples, coloured by site. On the right, the fraction of samples in which both *Prevotella* and either *Bacteroides* or *Phocaeicola* are present (relative abundance ≥1e-04) is shown across sites, indicating that these genera co-exist in most samples.

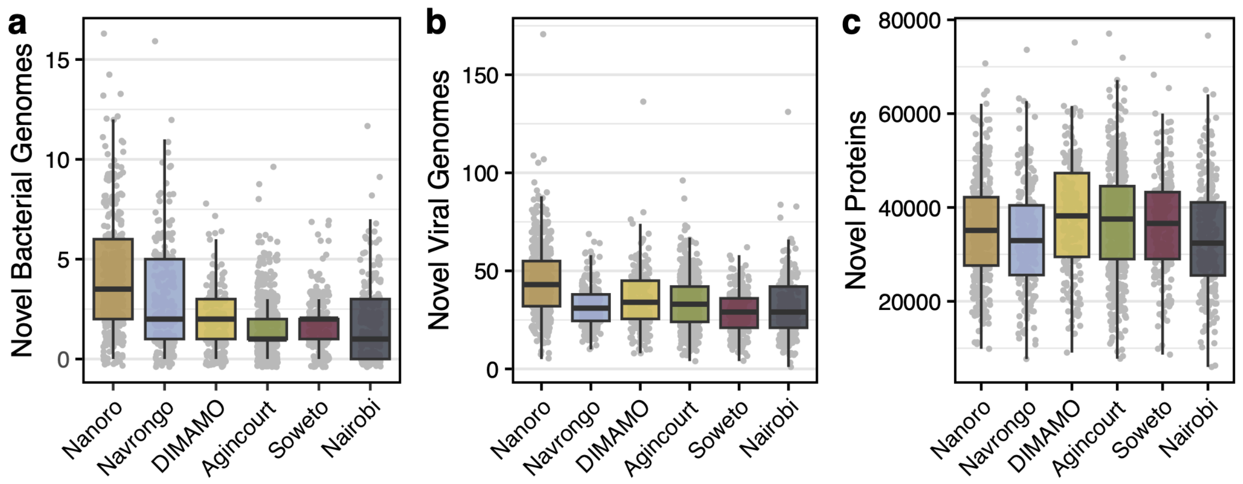


**Supplementary Figure 6. Novel microbial features per sample.** Number of novel **a)** prokaryotic genomes relative to the UHGG, **c)** viral genomes relative to the MGV, and **d)** prokaryotic proteins relative to the UHGP95 present in each sample. Points indicate the number of genome or protein clusters present per sample that are not found in respective feature databases. For all boxplots, boxes denote the interquartile range (IQR) with the median as a thick black line and the whiskers extending up to the most extreme points within 1.5-fold IQR.

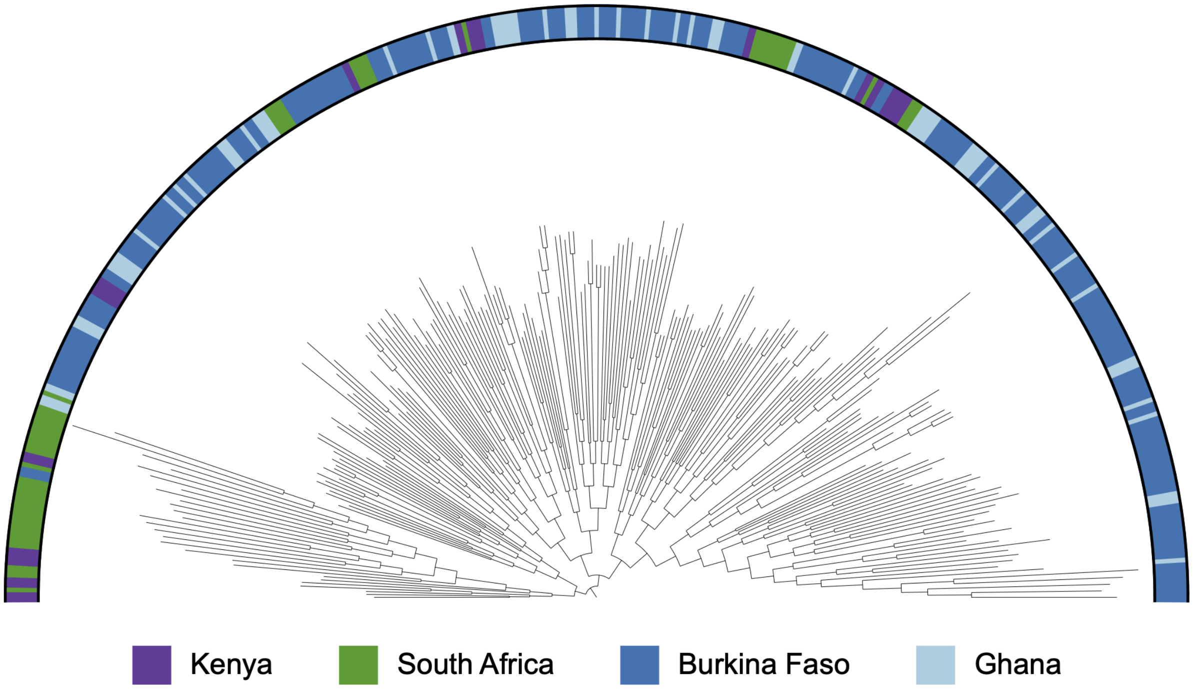


**Supplementary Figure 7. *Treponema succinifaciens* phylogeography in AWI-Gen samples.** Midpoint-rooted phylogenetic tree of 249 *T. succinifaciens* metagenome-assembled genomes generated from this study. Colour coding in the outside ring indicates the country of origin.

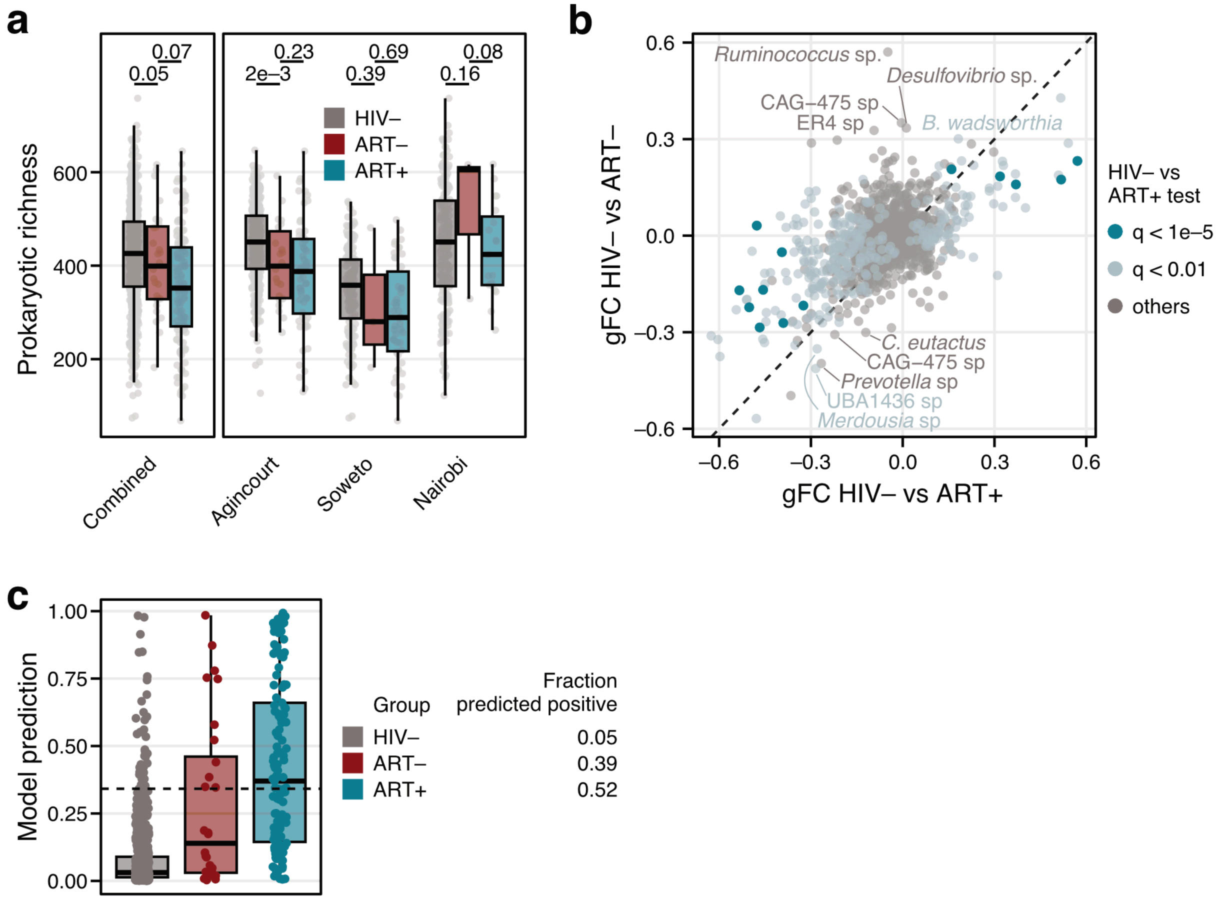


**Supplementary Figure 8. PLWH without ART exhibit microbiome changes similar to PLWH with ART. a**) Prokaryotic richness (number of prokaryotic species present at ≥1e-04% abundance) by HIV status, antiretroviral therapy, and site. Points represent individual samples. Differences in alpha diversity for each individual site were tested with ANOVA and for all sites combined with a linear mixed effect model accounting for site as a random effect. **b)** Generalized fold change (gFC) for all species between seronegative (HIV-) and PLWH with ART compared to the gFC between HIV- and PLWH without ART. Species are coloured by their q-value in the HIV- vs PLWH, ART+ comparison (see Figure 6 in the main text as well) and species with an absolute gFC ≥ 0.3 in the HIV- vs PLWH, ART- comparison (that do not exhibit a gFC ≥ 0.3 in the HIV- vs PLWH, ART+ comparison) are annotated (*C* = *Coprococcus*, *B* = *Bilophila*). **c)** Prediction from the machine learning model trained on all data by HIV status. The model was trained only on data from HIV- and PLWH, ART+ participants and applied to the PLWH, ART- participants as an external dataset. The fraction of samples predicted to be positive at the cutoff (horizontal dashed black line) corresponding to a 5% internal false positive rate on the HIV- samples are annotated on the right. For all boxplots, boxes denote the interquartile range (IQR) with the median as a thick black line and the whiskers extending up to the most extreme points within 1.5-fold IQR.
